# Interneuron activity-structural plasticity association is driven by context-dependent sensory experience

**DOI:** 10.1101/2021.01.28.428612

**Authors:** Soham Saha, John Hongyu Meng, Hermann Riecke, Georgios Agoranos, Kurt A. Sailor, Pierre-Marie Lledo

## Abstract

Neuronal dendritic spine dynamics provide a plasticity mechanism for altering brain circuit connectivity to integrate new information for learning and memory. Previous *in vivo* studies in the olfactory bulb (OB) showed that regional increases in activity caused localized spine stability, at a population level, yet how activity affects spine dynamics at an individual neuron level remains unknown. In this study, we tracked *in vivo* the correlation between an individual neuron’s activity and its dendritic spine dynamics of OB granule cell (GC) interneurons. Odor experience caused a consistent correlation between individual GC activity and spine stability. Dissecting the components of the OB circuit showed that increased principal cell (MC) activity was sufficient to drive this correlation, whereas cell-autonomously driven GC activity had no effect. A mathematical model was able to replicate the GC activity-spine stability correlation and showed MC output having improved odor discriminability while retaining odor memory. These results reveal that GC spine plasticity provides a sufficient network mechanism to decorrelate odors and maintain a memory trace.

## Introduction

Synaptic plasticity is the predominant mechanism of learning and memory in neural networks ^1–3^. During cortical development extensive structural plasticity, driven by coincident activity of neuronal partners, causes the formation and refinement of dendritic spines ^4, 5^. This form of structural plasticity also occurs in the adult brain, albeit at a much lower level; instead, different mechanisms of potentiating or depressing synaptic strengths dominate^6^.

Nonetheless, structural plasticity in the adult brain was shown to be modulated by sensory changes and new spine formation has been correlated with learning in the cerebral cortex ^7, 8^ and olfactory bulb ^9^ of adult mice.

Granule cells (GCs), the predominant interneuron of the olfactory bulb (OB), undergo persistently elevated structural plasticity throughout life, with approximately 20% of each GC’s spines turning over daily^10^. GCs inhibit the OB principal neurons, mitral and tufted cells (MCs), through reciprocal dendro-dendritic synapses ^11^. MC glutamate release onto a GC causes reciprocal GABA release from the GC and signal propagation along the GC dendrite recruits additional spines to laterally inhibit other MCs, shaping OB output to the olfactory cortex ^12^. In the sparsely connected OB circuit, structural plasticity may provide an efficient mechanism, in addition to classical synaptic plasticity, for the network to adjust for processing new complex sensory inputs that are characteristic of the high dimensional space of odorants ^13^.

Perceptual learning in the OB is an inherent form of learning where odor experience causes increased discriminability of similar experienced odor ^14^. The circuit mechanism of this learning is still poorly understood, but evidence suggests it to be driven by GC modulation of MC output ^15, 16^. Previous studies demonstrated that continuous odor exposure stabilized GC spines in active OB regions ^17^. Furthermore, activity-dependent glutamate and BDNF release from MCs was found to regulate GC spine head filopodia motility *in vitro* at short time-scales ^18^. However, it is yet unknown how individual neuron activity regulates GC structural plasticity within the timespan of functional spine development.

To determine the potential impact activity has in driving spine dynamics, we developed an awake structural *in vivo* imaging paradigm to track GC spine dynamics in response to odor experience. We further manipulated activity within the OB circuit to determine which components are essential for driving this plasticity. Finally, we established a computational model utilizing spine transition states to replicate the *in vivo* spine stability-activity correlation. Despite spines being highly dynamic, the model network demonstrated enhanced odor discriminability, while having the ability to retain memory.

## Results

### Awake *in vivo* 2-photon imaging of sparsely labelled GC neurons and reconstructing the spines

To understand the relationship between neuronal activity and dendritic spine dynamics at a single-cell resolution, we employed a viral labelling method for simultaneously detecting neuronal activity and spine dynamics of GC dendrites in awake animals. Sparse labelling of GCs was accomplished by injecting lentivirus expressing tdTomato (tdTom) and Cre-recombinase into the rostral migratory stream (RMS) to label neuroblasts ^19^ (Fig. 1a), after 3 weeks, adeno-associated virus expressing floxed genetically encoded calcium indicator (GCaMP6f) was injected in the OB and a cranial window over the OB was performed as described previously ^10^. Numerous studies tracking GC spine dynamics were exclusively performed in anaesthetized mice ^10, 17, 20^, however, GCs are highly influenced by anesthesia, causing an almost complete loss of spontaneous activity while strongly reduced odor responses ^21^. Therefore, we designed an awake protocol for imaging neuronal structure with sufficient resolution to track GC dendritic spines (Fig. 1b, Extended Data Fig. 1). The tdTom signal also provided a stable channel for detecting and removing movement artefact frames of the GCaMP signal (Extended Data Fig. 2). The details of the pre-processing steps for analysis of neuronal activity are described in the *Methods*.

**Figure 1:**
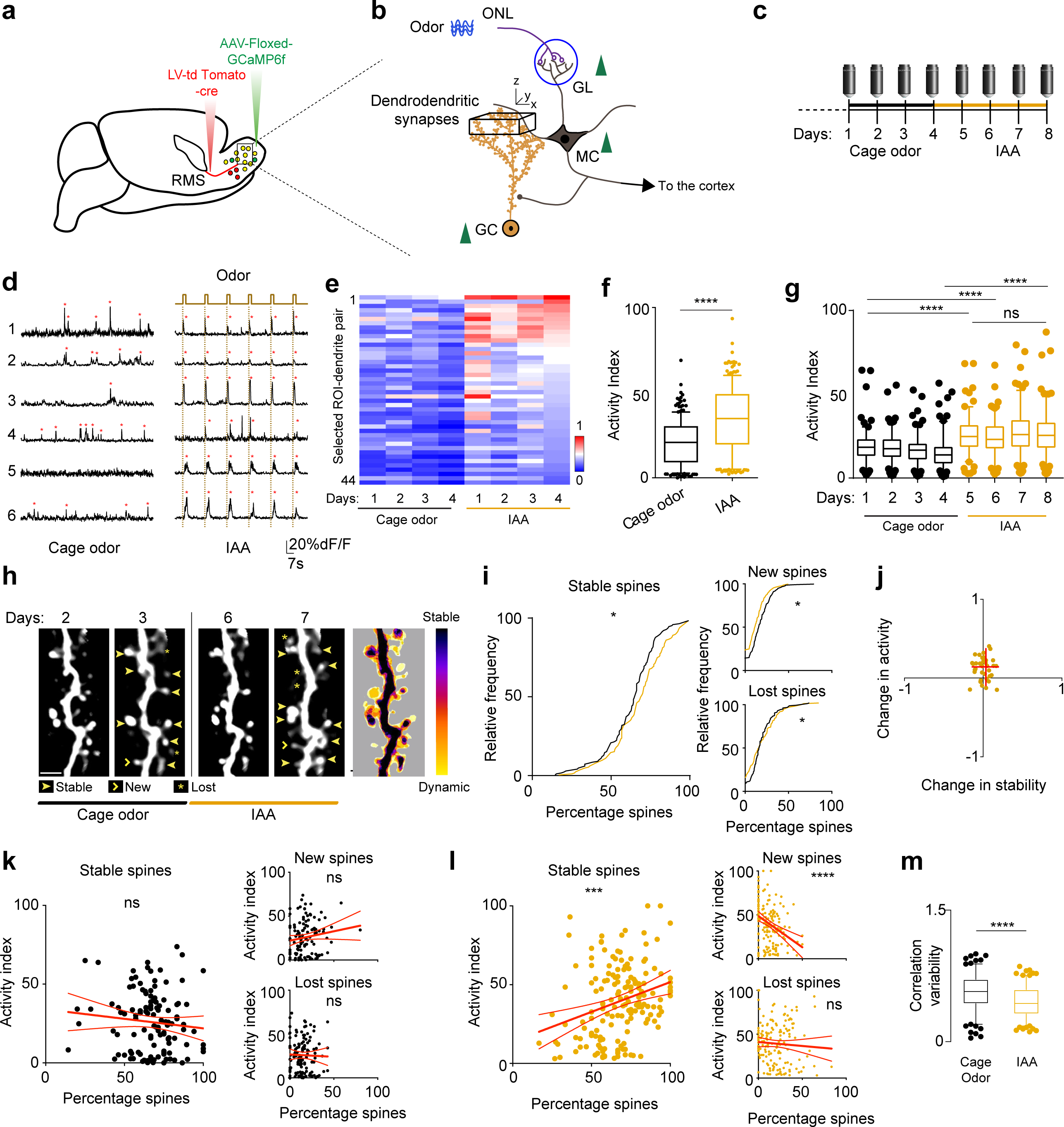
GC apical spine stability is highly correlated with continuous IAA-driven activity. a. Schematic representation of the viral labelling strategy to sparsely label GCs. Adult born neuroblast structure is labelled with lentivirus expressing tdTomato and Cre recombinase that is injected into the rostral migratory stream (RMS). The cells migrate to the OB, within three weeks differentiate into GCs, and AAV expressing floxed-stop GCaMP6f is injected to label tdTomato/Cre-expressing GCs for detecting calcium activity. b. OB circuit with mitral cells (MCs) receiving direct input from the GCs through dendro-dendritic synapses. Green arrows indicate increased activity of the cells with IAA (apical part of GC dendrites). The black cube in the GC apical dendrites demarcates the chronic *in-vivo* volume for structural 2-photon imaging and a single z-plane within this volume was used for measuring calcium activity. c. Experimental timeline for the *in vivo* imaging protocol. For the first 4 days, the mice were daily imaged in the ‘cage-odor’ condition (solid black line) following which a new odor (IAA) was placed in the cage for 4 days (solid gold line) and the mice were imaged daily. d. Example calcium traces of selected GCs in cage odor (spontaneous activity) and IAA (IAA; odor evoked activity) imaging trials with the same cells’ traces represented for each condition. IAA odor delivery period is indicated at the top. e. Heat map showing the neuronal activity (area under the curve-AUC, min-max normalized across the dataset) averaged per day across different experiments. The ROIs-dendrite pairs are selected based on the co-expression of tdTomato and GCaMP6f (44 dendrite-ROI pairs, 8 animals). f. Boxplots representing the mean of min-max normalized AUC (activity index) of all the ROI-dendrite pairs for the experimental conditions: cage odor and IAA. g. Boxplots representing the neuronal activity (activity index) in selected ROI-dendrite pairs (apical region of GCs) across days for different experiments (black: cage odor background, gold: IAA). h. Two-photon projected images from days 2 to 3 and 6 to 7, corresponding to cage odor and IAA, respectively. New, stable and lost spines for a 1-day interval as labeled in the figure. Heatmap binary overlay of days 1-8 of the same dendritic segment. Scale bar = 5 um. i. Cumulative frequency plot of stable, new and lost spines in cage odor background (black) and IAA (gold) conditions. j. Quadrant plot of the change in mean activity index versus the change in mean percentage of stable spines between cage odor and IAA conditions. Red cross indicates the standard deviation of the variables around the mean. k. Scatter plots of neuronal activity (activity index) versus percentage of stable, new and lost spines in individual dendrite-ROI pairs in cage odor with linear regression fit. l. Scatter plot of neuronal activity (activity index) versus percentage of stable, new and lost spines in individual dendrite-ROI pairs in IAA conditions with linear regression fit. m. Boxplot of the correlation variability of the individual dendrite-ROIs in scatter plot (panel k and l) from their mean for cage odor background (black) and IAA (gold).

### Odor experience drives apical dendro-dendritic spine stability

The relationship between GC neuronal activity and its apical dendritic spine stability was explored with animals in normal cage conditions for 4 days with daily imaging, followed by 4 days of exposure to the odorant isoamyl acetate (IAA) in the cage with daily imaging (Fig. 1c). On the microscope stage, a mask provided the mouse with continuous air that first passed through each animal’s matched cage bedding. This “cage odor” was used for all experiments to mimic the cage odor environment to avoid off-target effects on both activity and structural plasticity since there was a significant difference in GC activity between pure air and cage odor presentation (see Extended Data Figure 3f; p = 0.012).

The tdTom signal was imaged daily as a z-series volume to reconstruct apical dendritic spine segments for tracking spine dynamics. A single, defined z-plane within the center of this volume was imaged daily as a time-series at high speed (15 Hz) for sufficient temporal resolution to record GCaMP6f-labeled dendritic activity (Extended Data Fig. 3a, b). In the “cage odor” condition during days 1-4, spontaneous activity was recorded for activity baseline, whereas in the “IAA” condition, during days 5-8, odor evoked activity was recorded with 2s of IAA delivery per trial.

Activity in the GC dendrites increased significantly with IAA exposure (Fig. 1d-f; Table 5; p < 0.0001, n = 8 animals, 51 dendrite-ROI pairs), which was sustained over subsequent days (Fig. 1g), showing no significant habituation effect. The activity of neighboring ROIs along the same dendritic branch was highly correlated (Extended Data Fig. 3d-e; F-test, p = 0.797), which allowed associating structural dynamics of individual spines with their common dendritic segment activity. Exposure to IAA increased the overall stability of spines (Fig. 1h-i; p = 0.02; n = 8 animals, 51 dendrite-ROI pairs; Extended Data Table 5), while reducing the population of new and lost spines (Fig. 1i; new p = 0.018; lost p = 0.04; for detailed statistics of all figures see Extended Data table 5).

To explore the effect of odor experience on spines, the mean of activity and spine stability with IAA for each segment, minus the mean of activity and spine stability with baseline cage odor conditions, was plotted (Fig. 1j). The upper right quadrant shift suggested a cumulative positive activity and spine stability change with IAA. The effect of odor experience on spine dynamics was assessed in more detail using a linear regression fit between the activity across days for each segment and the corresponding spine classification. With cage odor, there was no significant correlation with activity in new, lost or stable spines (Fig. 1k; stable, p = 0.269; new, p = 0.11; lost, p = 0.756). IAA, however, caused a significant positive correlation between dendritic activity and spine stability (Fig. 1l; Extended Data Fig. 3g; p = 0.0002) and a significant negative correlation with new spine formation (Fig. 1l; p < 0.0001; Extended Data Fig. 3h) with no significant correlation in lost spines (Fig. 1l; p = 0.414; Extended Data Fig. 3i).

To determine the variability of correlation across days, the spread of the dataset was measured by taking the distance of all individual daily activity-stability points for each dendrite from their respective means across the experimental paradigms all days, segregating between cage odor and IAA (Fig. 1m). The variability of the correlation reduced significantly with IAA, as compared to cage odor (Fig. 1m; cage odor: 0.56 ± 0.20, IAA: 0.45 ± 0.18; p < 0.0001), indicating odor experience leads to higher spine stability across days on individual spines/segments. Overall, these data suggest that odor experience drives sustained spine stability and is sufficient to reduce the dynamic spine pool, driving an activity-spine stability correlation, while reducing inter-day differences in this correlation.

### Odor-induced activity does not drive proximal spine stability

In contrast to GC apical spines which make reciprocal dendro-dendritic synapses, proximal spines, those that are on the primary apical dendrite, immediately adjacent and superficial to the soma, have classical spines (Extended Data Fig. 4a-b), predominantly receiving top-down input. We explored whether the same activity-spine stability relationship rule could be extended to proximal spines. The activity and spine dynamics at the proximal dendrite compartment were measured using an identical odor experience paradigm as in Fig. 1 (Fig. 2a-b). IAA exposure caused a robust increase in neuronal activity (Fig. 2c-e; p <0.0001; n = 4 animals, 55 dendrite-ROI pairs) while there was also a significant increase in overall spine stability (Fig. 2f-g; p < 0.0001) coupled with a significant decrease in the fraction of new spines (Fig. 2g; p < 0.0001), and no change in lost spines (Fig. 2g; p = 0.095).

**Figure 2:**
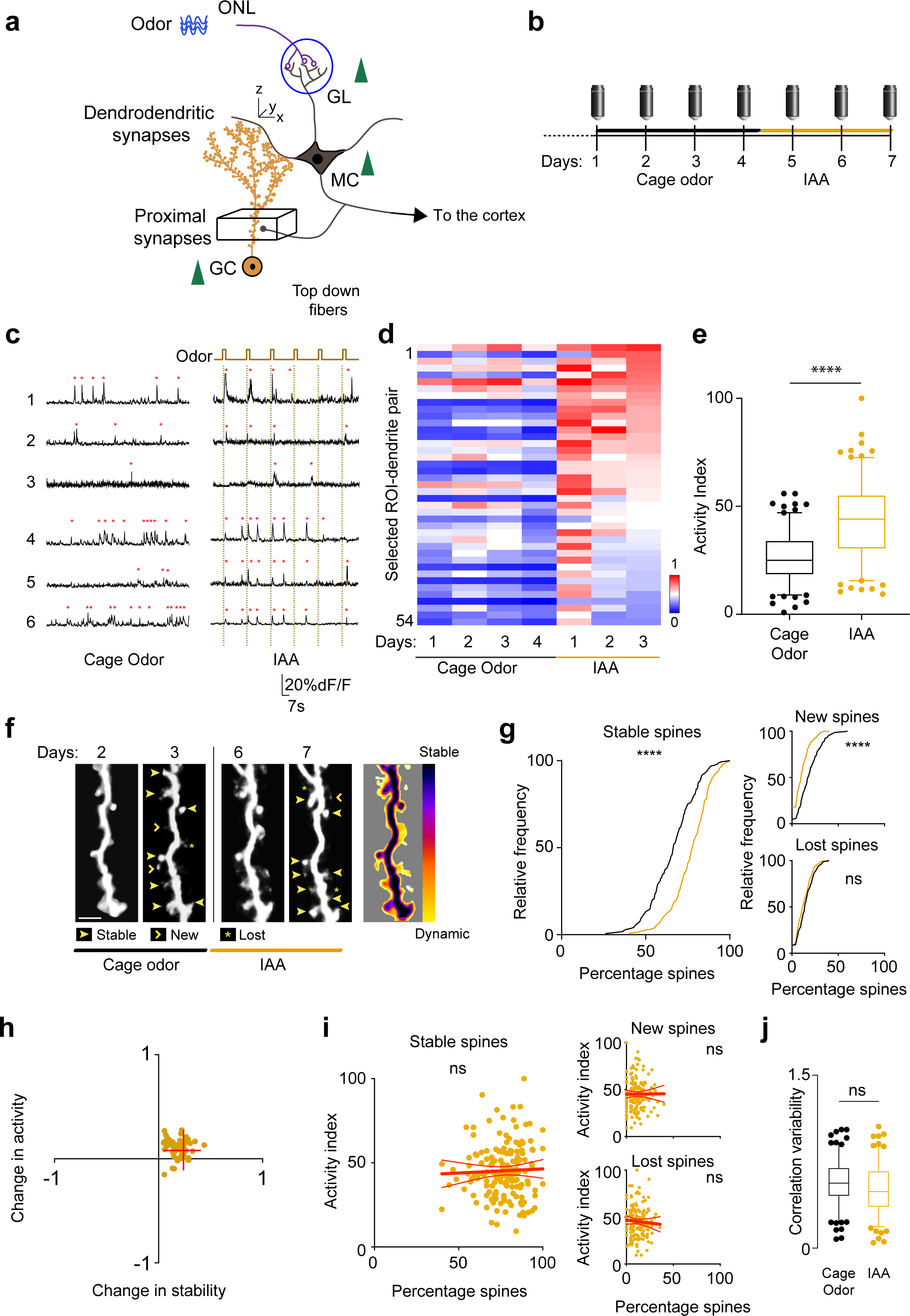
GC proximal spine stability is not correlated with its individual activity. a. OB circuit as outlined in figure 1. The black cube in the GC proximal dendrites demarcates the chronic *in-vivo* volume for structural 2-photon imaging and a plane within this volume was used for measuring calcium activity. b. Experimental timeline as outlined in Fig.1. The mice were 2-photon imaged in the ‘cage-odor’ condition (solid black line) for 4 days following which a new odor was placed in the cage for 3 days (solid gold line) and the mice were imaged in the ‘IAA’ condition. c. Example calcium traces of selected GCs in cage odor (spontaneous activity) and IAA (odor evoked activity) imaging trials with the same cells’ traces represented for each condition. Odor delivery period is indicated at the top. d. Heat map showing the neuronal activity (AUC, min-max normalized across the dataset) averaged per day across different experiments (54 dendrite-ROI pairs, 4 animals). e. Boxplots representing the neuronal activity (AUC, min-max normalized) in selected ROI-dendrite pairs (proximal region of GCs) across days for different experiments (black: cage odor background, gold: continuous IAA). f. 2-photon projected images from day 2 to 3 and 6 to 7, corresponding to cage odor and continuous IAA, respectively. New, stable and lost spines for a 1-day interval as indicated in the figure. Heatmap binary overlay of all the days of the same dendritic filament with colder colors indicating more stable spines. Scale bar = 5um. g. Cumulative frequency plot of stable (left), new and lost spines (right) in cage odor background (black) and continuous IAA (gold) showing significant increase in the percentage stable spines with continuous IAA. The new spines are significantly reduced (p < 0.0001) which no change in lost spines are seen. h. A quadrant plot of the change in mean activity index versus the change in mean percentage of stable spines between cage odor background and IAA for each ROI. The red cross indicates Mean ± SD of both the parameters. i. Scatter plot of neuronal activity (activity index) versus percentage of stable spines (left), new and lost spines (small, right) in individual dendrite-ROI pairs in continuous IAA with linear regression fit. j. Boxplot of the correlation variability of the individual dendrite-ROIs from their mean for cage odor background (black) and continuous IAA (gold).

Although, both mean activity and mean spine stability were increased during IAA exposure (Fig. 2h), there was no significant correlation between individual segment activity and new, lost or stable spines (Fig. 2i; *new:* p = 0.951; *lost*: p = 0.447; *stable*: p = 0.647; Extended Data Fig. 4c-e; Extended Data table 3). In addition, the decrease in the correlation variability across days with IAA was not significant from cage odor background (Fig. 2j *cage odor*: 0.57 ± 0.19, *IAA*: 0.50 ± 0.22; p = 0.085). Thus, proximal spines have limited or no correlation with their structural plasticity in response to direct sensory stimulation.

### Short-term odor exposure is incapable of driving spine stability

After determining that apical spines have a unique odor-induced activity-stability relationship, we explored the effect of the duration of odor experience on spines. Instead of continuous IAA exposure in their cages, mice were presented with IAA for 40, 2s presentations per day during the imaging sessions (Fig. 3a-b; ‘short-term IAA’). Evoked IAA short-term exposure caused a significant increase in activity across days (Fig. 3c-e; p < 0.0001; 5 animals; 42 dendrite-ROI pairs).

**Figure 3:**
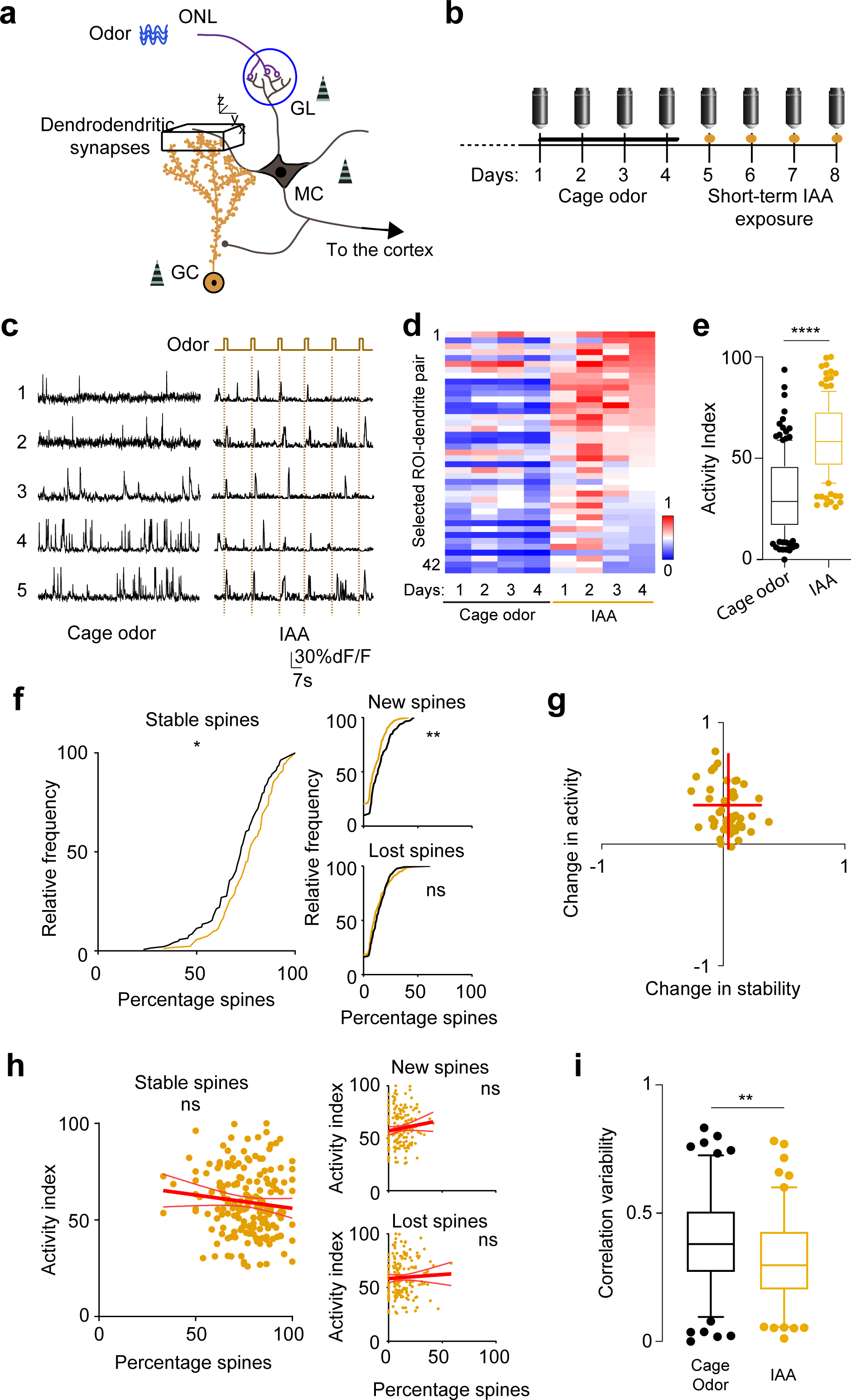
Short term exposure to odor does not lead to correlated increase in activity and spine stability in apical GC dendrites. a. Outline of the OB circuit. Black cube in the GC apical dendrites demarcates the chronic *in-vivo* volume for structural 2-photon imaging and a plane within this volume was used for measuring calcium activity. b. Experimental timeline as outlined in Figure 1 & 2. The mice were 2-photon imaged in the ‘cage-odor’ condition (solid black line) for 4 days followed by short-term exposure to the same odorant as in Figure 1 passively for 15 mins for the next 4 days (gold dots on the timeline). c. Example calcium traces of selected GCs in cage odor (spontaneous activity) and short-term passive IAA (odor evoked activity) imaging trials with the same cells’ traces represented for each condition. Odor delivery period is indicated at the top. d. Heat map showing the neuronal activity (AUC, min-max normalized across the dataset) averaged per day across different experiments (42 dendrite-ROI pairs, 4 animals). e. Boxplots representing the mean of min-max normalized AUC of all the ROI-dendrite pairs for the experimental conditions: cage odor and short-term passive IAA. f. Cumulative frequency plot of stable (left), new and lost spines (right) in cage odor background (black) and short-term IAA (gold) showing significant increase in the percentage stable spines with short-term IAA. g. A quadrant plot of the change in mean activity index versus the change in mean percentage of stable spines between cage odor background and short-term IAA for each ROI. The red cross indicates Mean ± SD of both the parameters. h. Scatter plot of neuronal activity (activity index) versus percentage of stable spines (left), new and lost spines (small, right) in individual dendrite-ROI pairs in short-term IAA with linear regression fit. i. Boxplot of the correlation variability of the individual dendrite-ROIs from their mean for cage odor background (black) and short-term IAA (gold).

There was a slight increase in spine stability (Fig. 3f, p = 0.048), with significantly fewer new spines in ‘short-term IAA’ compared to ‘cage odor’ (Fig. 3f; p = 0.003) and no significant change in lost spines (Fig. 3f; p = 0.299). The displacement of the change in activity-stability scatter was characterized by an increase in activity (Fig. 3g). There was no significant correlation between individual dendritic activity and the percentage of stable, new or lost spines (Fig. 3h; *new:* p = 0.159; *lost*: p = 0.529; *stable*: p = 0.153). Interestingly, the correlation variability across days showed a significant decrease between cage odor and short-term IAA (Fig. 3i; *cage odor*: 0.38 ± 0.18, *short-term IAA*: 0.28 ± 0.16; p = 0.005), but was significantly less compared to continuous IAA (p = 0.032). These results indicate that the experience of transient odor exposure is not sufficient to drive the GC activity-spine stability correlation.

### Activity-dependent spine stability is driven by sensory-input

GC apical spines are stabilized by long-term odor exposure, but is sensory-driven activity necessary for maintaining spine stability? Sensory deprivation by unilateral naris occlusion was used to decrease network activity, but spare spontaneous activity, to explore the effect on GC apical spine stability ^22^. In addition to the protocol in Fig. 1, the mice underwent naris occlusion after the IAA exposure period and GC apical dendrites were imaged daily in all conditions (Fig. 4a-b).

**Figure 4:**
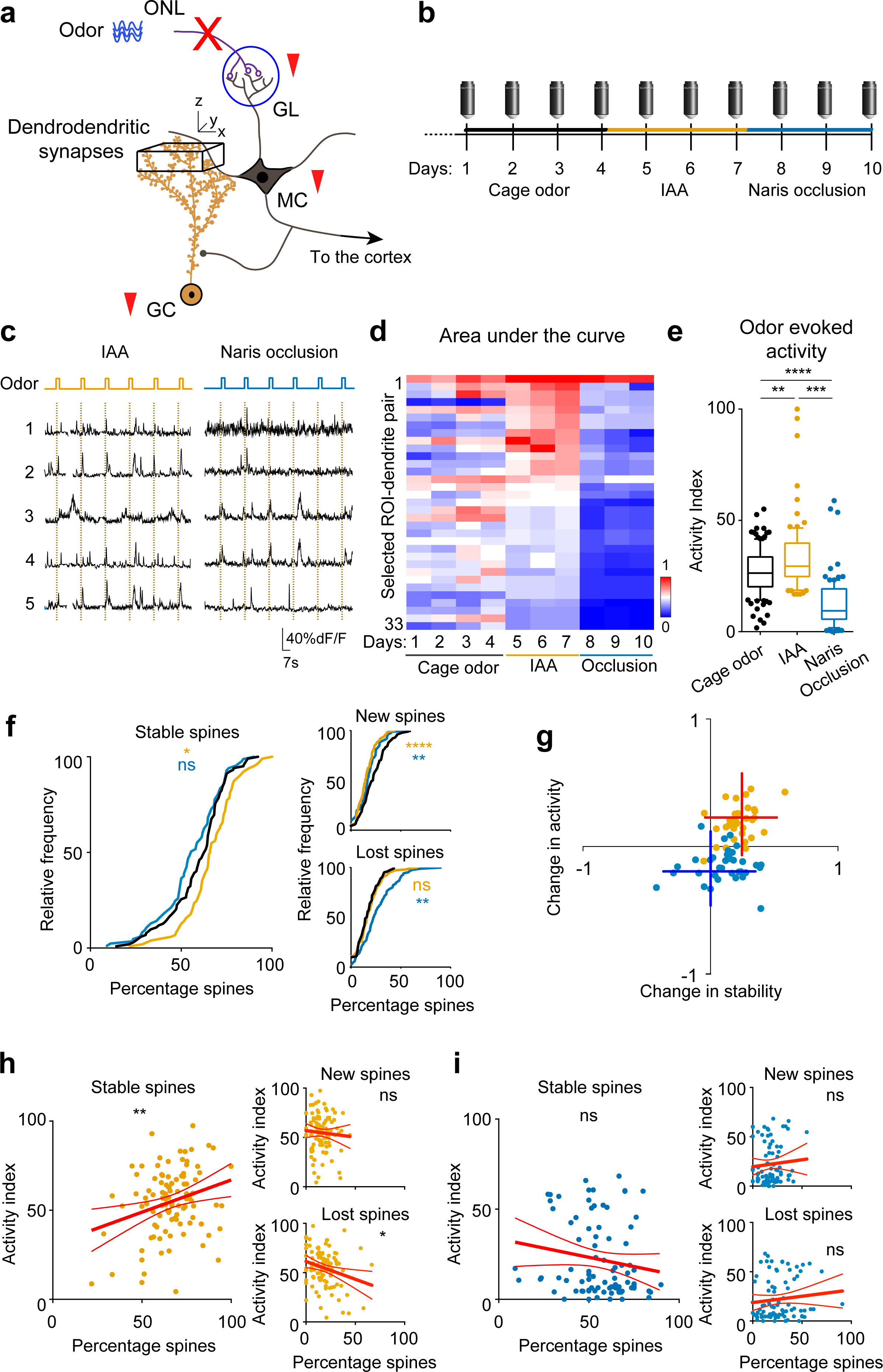
Sensory deprivation disrupts the positive relationship of neuronal activity and spine stability established by continuous sensory exposure. a. OB circuit as outlined previously. Black cube in the GC apical dendrites demarcates the chronic *in-vivo* volume for structural 2-photon imaging and a plane within this volume was used for measuring calcium activity. Unilateral naris occlusion was used to block sensory input from the olfactory nerve layer (ONL) to the OB. b. Experimental timeline as outlined in Figure 1. The mice were 2-photon imaged in the ‘cage-odor’ condition (solid black line) for 4 days following where IAA was placed in the cage for 3 days (solid gold line) and the mice were imaged. Afterwards, the mice were imaged with unilateral ‘naris occlusion’ condition for the last 3 days (solid blue line). c. Example calcium traces of selected GCs in continuous IAA (odor evoked activity) and naris occlusion (odor evoked activity) imaging trials with the same cells’ traces represented for each condition. Odor delivery period is indicated at the top. d. Heat map showing the neuronal activity (AUC, min-max normalized across the dataset) averaged per day across different experiments (33 dendrite-ROI pairs, 6 animals). e. Boxplots representing the min-max normalized AUC of all the ROI-dendrite pairs for the experimental conditions: cage odor, IAA and naris occlusion. f. Cumulative frequency plot of stable (left), new and lost spines (small, right) in the cage odor background (black), continuous IAA (gold) and naris occlusion (blue). g. A quadrant plot of the change in mean activity index versus the change in mean percentage of stable spines between cage odor background and continuous IAA (in gold); and naris occlusion and cage odor background (in blue) for each ROI. h. Scatter plot of neuronal activity (activity index) versus percentage of stable (left), new and lost spines (small, right) in continuous IAA with linear regression fit. i. Scatter plot of neuronal activity (activity index) versus percentage of stable (left), new and lost spines (small, right) in naris occlusion with linear regression fit.

Naris occlusion caused a significant decrease in apical GC dendritic activity as compared to cage odor spontaneous activity and IAA-evoked activity (Fig. 4c-e; *cage odor vs IAA*: p = 0.002, *cage odor vs naris*: p < 0.0001, *IAA* vs *naris*: p < 0.0001; 7 animals; 53 dendrite-ROI pairs; Extended Data. Fig. 5a-c). As in Fig.1, continuous IAA exposure led to a significant increase in spine stability (Fig. 4f; p = 0.039). However, compared to the cage condition, no significant change in stability was observed with naris occlusion (Fig. 4f; p = 0.135), while it significantly decreased the percentage of new (Fig. 4f; p = 0.009) and lost spines (Fig. 4f; p = 0.001).

As compared to spontaneous cage odor activity, naris occlusion reduced the mean activity without strongly affecting stability. During the IAA exposure period, both activity and stability were enhanced compared to the cage-odor period (Fig. 4g; Δstability (IAA) = 0.219 ± 127; Δactivity (IAA) = 0.190 ± 134; Δstability (naris) = 0.102 ± 191; Δactivity (naris) = -0.152 ± 126). IAA exposure pushed the activity-stability relationship to a significant positive correlation (Fig. 4h; p = 0.006; Extended Data Fig. 5d) with no change in new spines (Fig. 4h; p = 0.504; Extended Data Fig. 5e), but a slightly negative correlation for the lost spines (Fig. 4h; p = 0.012; Extended Data Fig. 5f).

During naris occlusion there was no significant correlation between activity and spine stability (Fig. 4i; Extended Data. Fig. 5g-i). Consistent with previous observations, the correlation variability across days was significantly less in IAA compared to cage odor and naris occlusion (Extended Data Fig. 5c). These results suggest that odor exposure is required for establishing stable connectivity between GCs and the principal cells with the disruption of sensory input leading to their destabilization.

We co-labelled the GCs with tdTom and a lentivirus expressing GFP fused to PSD-95 (Extended Data Fig 6a-b) ^17^, to track the proportion of “functional spines” (tdTom^+^/PSD-95^+^) and potential “proto-spines” (tdTom^+^/PSD-95^-^) under cage odor, IAA, and naris occlusion conditions. We observed that not all spines express PSD-95 (Extended Data Fig 6d). In addition, with naris occlusion, only the density of PSD-95^+^ spines were reduced (Extended Data Fig 6c). The PSD-95^+^ spine fraction was insignificantly increased with IAA and similarly decreased with sensory deprivation (Extended Data Fig 6d). Although PSD-95^+^ and PSD-95^-^ spines have increased stability with IAA, and reduced stability with naris occlusion (Extended Data Fig 6h; shown by the centroid distance from the origin), IAA does not lead to correlated stability in PSD-95^+^ and PSD-95^-^ spines (Extended Data Fig 6e-g; Cage odor: p = 0.09 (stable), p = 0.002 (new); IAA: p = 0.157 (stable), p = 0.224 (new); naris occlusion: p = 0.01 (stable), p = 0.08 (new)). However, the rates of PSD-95^+^ and PSD-95^-^ spine loss in both IAA and naris occlusion are highly correlated (Extended Data Fig 6e-g; Cage odor: p = 0.568 (lost); IAA: p = 0.0008 (lost); naris occlusion: p = 0.008 (lost)). Given these observations, we conclude that the rates of spine formation and spine consolidation are different in the case of olfactory sensory experience. These observations provide the rational of classifying spines into “functional” and “non-functional” spines in the computational model described later.

### GC silencing or activating does not alter the activity-stability relationship

To further dissect the OB components driving the spine activity-stability relationship, we utilized chemogenetic manipulation with Designer Receptor Exclusively Activated by Designer Drugs (DREADD) expressed in GCs to control their cell-autonomous activity ^23, 24^ (Fig. 5a; Extended Data Fig. 7a-b). GC activity was modulated by adding the DREADD ligand clozapine-N-oxide (CNO) into the cage drinking water (Fig. 5b). A separate cohort of mice were viral labeled with inhibitory (hM4Di) or excitatory (hM3Dq) DREADD to silence or activate GCs with CNO administration, respectively (Fig. 5a; Extended Data Fig. 7a).

**Figure 5:**
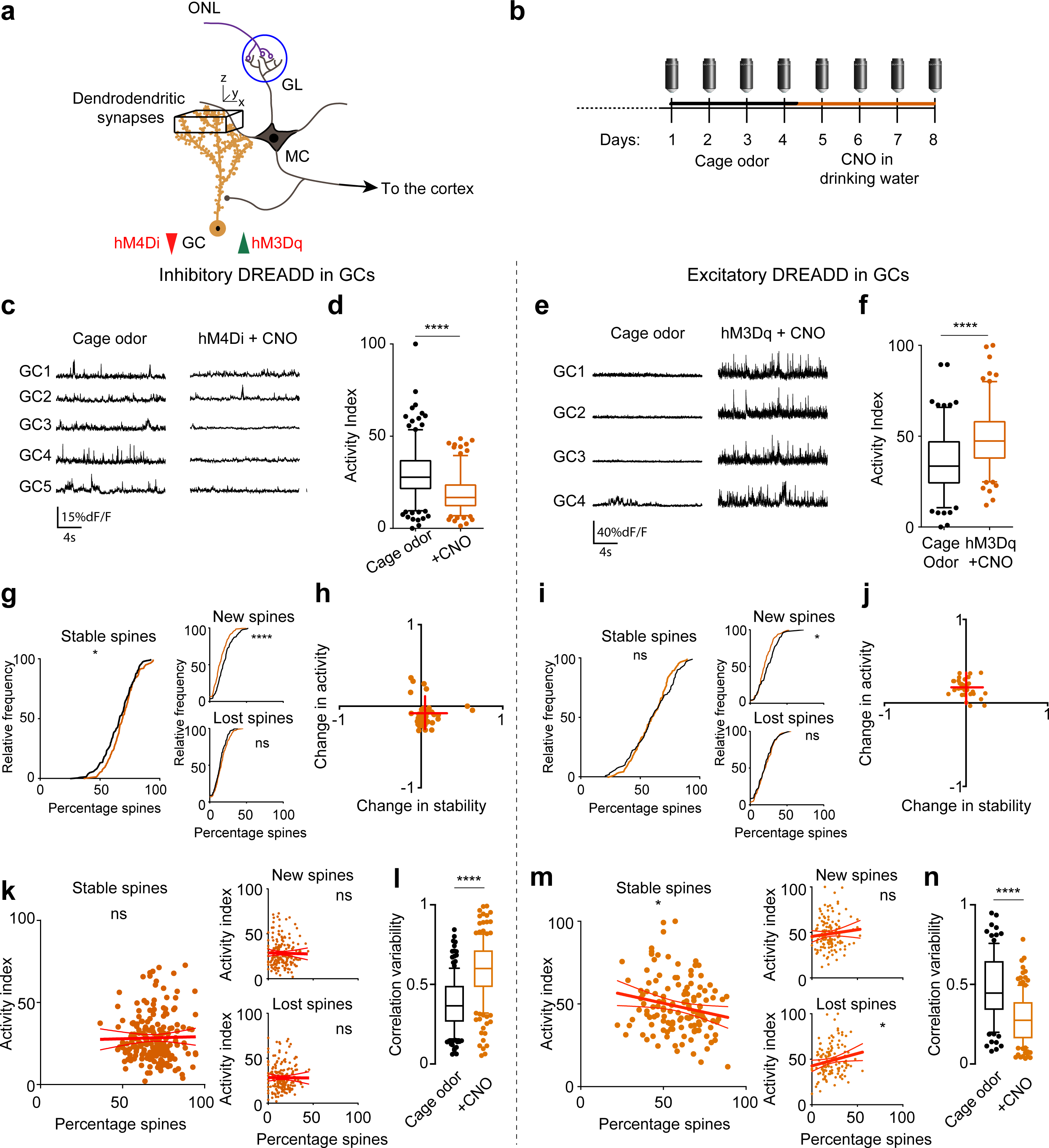
Cell autonomous suppression and activation of GC activity stabilizes spines but does not lead to correlated activity-spine stability relationship. a. OB circuit as outlined previously. Black cube in the GC apical dendrites demarcates the chronic *in vivo* volume for structural 2-photon imaging and a plane within this volume was used for measuring calcium activity. Red arrow indicates suppression and green arrow indicates activation. b. Experimental timeline as outlined previously. AAV mixture expressing floxed-stop GCaMP6f and floxed-hM4Di (inhibitory DREADD)/ floxed-hM3Dq (excitatory DREADD) is injected to label tdTomato/Cre-expressing GCs for detecting calcium activity and induce inhibition/activation of the GCs in a ligand (CNO)-dependent manner. The mice were 2-photon imaged in the ‘cage-odor’ condition (solid black line) for 4 days followed by introduction of CNO in the drinking water for the next 4 days (solid orange line). c. Example calcium traces of selected GCs in cage odor (spontaneous activity) and CNO-mediated inhibition (spontaneous activity) imaging trials with the same cells’ traces represented for each condition. d. Boxplots representing the mean of min-max normalized AUC of all the ROI-dendrite pairs for the experimental conditions: cage odor and CNO-mediated inhibition. e. Example calcium traces of selected GCs in cage odor (spontaneous activity) and CNO-mediated excitation (spontaneous activity) imaging trials with the same cells’ traces represented for each condition. f. Boxplots representing the mean of min-max normalized AUC of all the ROI-dendrite pairs for the experimental conditions: cage odor and CNO-mediated excitation. g. Cumulative frequency plot of stable, new and lost spines in the cage odor background (black) and CNO-mediated inhibition (orange). There is an increase in the stable spines and decrease in the new spines with CNO-mediated inhibition compared to the cage odor background. h. A quadrant plot of the change in mean activity index versus the change in mean percentage of stable spines between cage odor background and CNO-mediated inhibition for each ROI. The red crosses indicate Mean ± SD of both the parameters. i. Cumulative frequency plot of stable, new and spines in the cage odor background (black) and CNO-mediated excitation (orange). No significant change in the percentage stable spines with CNO-mediated excitation is observed compared to the cage odor background. j. A quadrant plot of the change in mean activity index versus the change in mean percentage of stable spines between cage odor background and CNO-mediated excitation for each ROI. The red crosses indicate Mean ± SD of both the parameters. k. Scatter plot of neuronal activity index versus percentage of stable spines (left), new and lost spines (small, right) in CNO-mediated inhibition with linear regression fit. l. Boxplot showing the correlation variability of the individual dendrite-ROI pairs in CNO-mediated inhibition is increased compared to the cage odor background. m. Scatter plot of neuronal activity index versus percentage of stable spines (left), new and lost spines (small, right) in CNO-mediated excitation with linear regression fit. n. Boxplot showing the correlation variability of the individual dendrite-ROI pairs in CNO-mediated excitation is decreased compared to the cage odor background.

GCs used for analysis were selected based on their multiple-expression of tdTom (for structure), GCaMP6f (for activity) and DREADD (hM4Di/hM3Dq-mCherry, as determined by using different 2-photon wavelengths for spectral subtraction (Extended Data Fig. 7a-b). GCs expressing inhibitory DREADD (hM4Di) had significantly suppressed activity with CNO application (Fig. 5c-d; Extended Data Fig. 7c; p < 0.0001; 5 animals; 49 dendrite-ROI pairs), while GCs expressing excitatory DREADD (hM3Dq) had significantly increased activity (Fig. 5e-f; Extended Data Fig. 8a-b; p < 0.0001; 4 animals; 36 dendrite-ROI pairs).

Spine stability was slightly, but significantly higher in GCs inhibited by hM4Di as compared to cage odor baseline dynamics (Fig. 5g; p = 0.04; Extended Data Fig. 7d) while excitation with hM3Dq did not show any significant change in spine stability (Fig. 5i; p = 0.184; Extended Data Fig. 8c). In both the cases, however, new spine formation was reduced significantly (Fig. 5g; *new* p < 0.0001; Fig. 5i; *new* p = 0.029; Extended Data Fig. 7e, 8d), while changes in spine loss were insignificant (Fig. 5g; *lost* p = 0.132; Fig. 5i; *lost* p = 0.818; Extended Data Fig. 7f, 8e).

Comparing continuous inhibition and excitation of GCs, there was a negative shift in the change in activity from baseline cage odor conditions in the hM4Di group (Fig. 5h) and an increase in activity in the hM3Dq group (Fig. 5j). Inhibition slightly enhanced the overall stability of the spines, while excitation had no significant influence, yet both groups had a slight reduction in the number of new spines. Inhibition of GCs caused no correlation between individual segment activity and the percentage of stable, new or lost spines (Fig. 5k; *new:* p = 0.803; *lost*: p = 0.927; *stable*: p = 0.773). A slight negative correlation in stability was found with GC excitation and conversely a slight positive correlation with lost spines while no correlation with new spines (Fig. 5l; *new:* p = 0.303; *lost*: p = 0.04; *stable*: p = 0.025). The correlation variability across days also increased with inhibition (Fig. 5l; hM4Di: *cage odor*-0.38 ± 0.19; *CNO* - 0.58 ± 0.19; p <0.0001), while decreased with excitation (Fig. 5n; hM3Dq: *cage odor*- 0.47 ± 0.21; *CNO* - 0.28 ± 0.16; p <0.0001). These data indicate that cell autonomous activation of GCs is not sufficient to drive the activity-stability correlation and, even more surprisingly, cell autonomous GC excitation appears to cause loss of spines.

### Persistent activation of MCs is required to drive the activity-stability relationship

A potential candidate for non-cell autonomous activity regulating spine stability is the primary input to GCs through MC activity. Recent *in vitro* evidence suggests that MC activity-dependent glutamate release regulates GC short-term spine motility^18^. We therefore targeted MCs by injecting AAV expressing floxed excitatory DREADD into the OB in Tbet-cre mice ^25^ (Fig. 6a). To mimic the continuous odor conditions for perceptual learning in Fig. 1, mice had baseline cage odor conditions with imaging followed by CNO in their cage drinking water with daily imaging (Fig. 6b).

**Figure 6:**
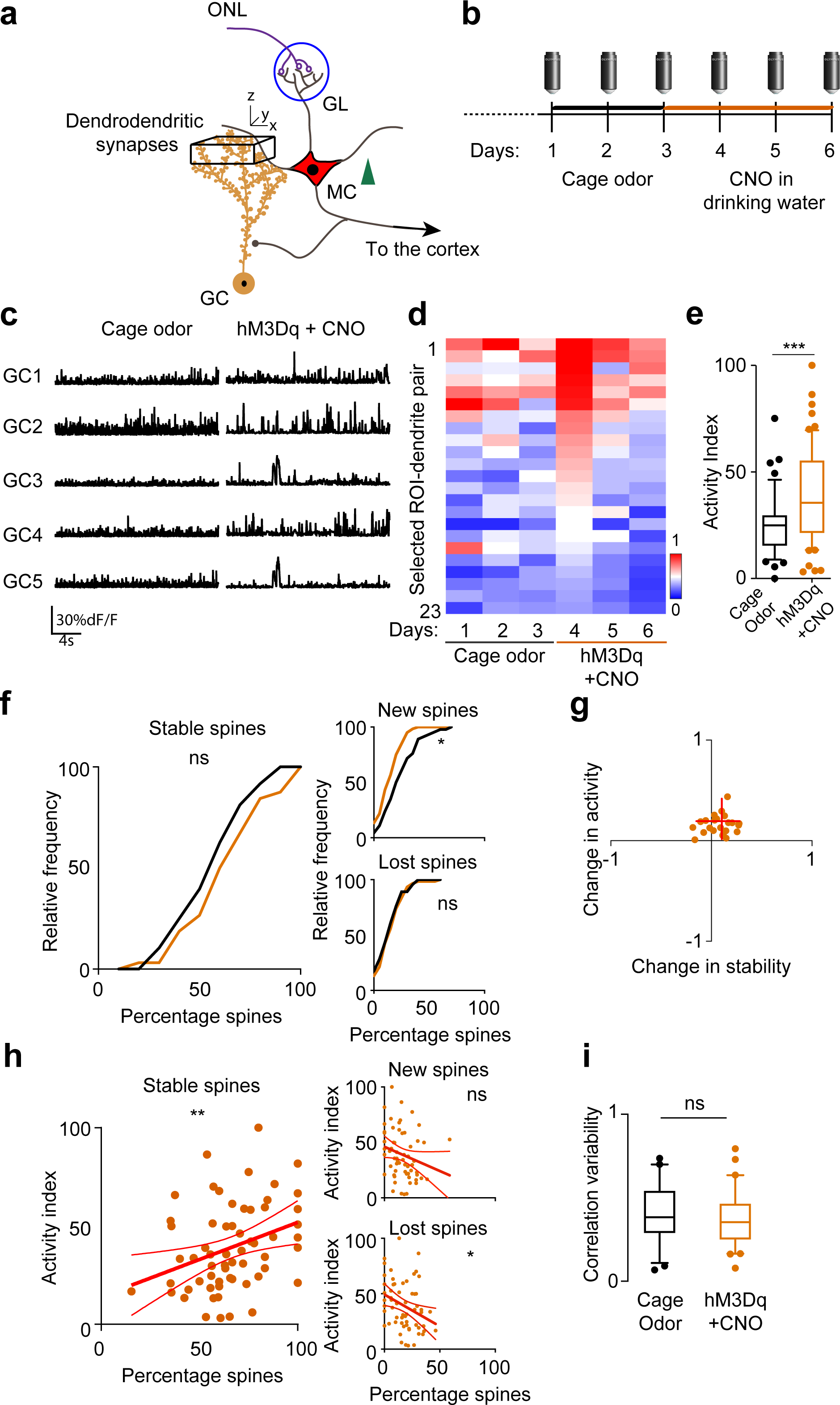
Chronic activation of MCs recapitulates the effect of odor-induced activity-dependent spine stabilization on GCs. a. OB circuit as outlined previously. Black cube in the GC apical dendrites demarcates the chronic in-vivo volume for structural 2-photon imaging and a plane within this volume was used for measuring calcium activity. b. Experimental timeline as outlined previously. Floxed-hM3Dq (excitatory DREADD) was injected in a Tbet-cre mouse to label the MCs specifically. The mice were 2-photon imaged in the ‘cage-odor’ condition (solid black line) for 3 days followed by introduction of CNO in the drinking water for the next 3 days (solid red line). c. Example calcium traces of selected GCs in cage odor (spontaneous activity) and CNO-mediated MC excitation (spontaneous activity) imaging trials with the same cells’ traces represented for each condition. d. Heat map showing the neuronal activity (AUC, min-max normalized across the dataset) averaged per day across different experiments (23 dendrite-ROI pairs, 4 animals). e. Boxplots representing the mean of min-max normalized AUC of all the ROI-dendrite pairs for the experimental conditions: cage odor and CNO-mediated MC excitation. f. Cumulative frequency plot of stable, new and lost spines in the cage odor background (black) and CNO-mediated MC excitation (orange). No significant change in the percentage stable spines with CNO-mediated MC excitation is observed compared to the cage odor background (KS test), while new spines significantly decreased. g. A quadrant plot of the mean change in activity index versus the mean change in percentage of stable spines between cage odor background and CNO-mediated MC excitation for each ROI. The red cross indicates Mean ± SD of both the parameters. h. Scatter plot of neuronal activity (activity index) versus percentage of stable (left panel), new and lost spines (small right panels) in CNO-mediated MC excitation (red) with linear regression fit. i. Boxplot of the correlation variability of the individual dendrite-ROIs in scatter plot from the mean scatterplot (panel I) for cage odor background (black) and CNO-mediated MC excitation (orange).

GC activity increased significantly with CNO when averaged across days (Fig. 6c-e; p = 0.0006; n = 4 animals, 23 dendrite-ROI pairs), with insignificant change on days 5 and 6 compared to day 1 (Extended Data Fig. 9a; p= 0.65, p = 0.88 respectively), and average spine stability increased, without reaching significance (Fig. 6f; p = 0.51). When compared across days, a significant increase in stability was seen on the 2^nd^ day of CNO application (Extended Data Fig. 9b; p = 0.034). A positive change in mean activity and spine stability with MC activation was also observed compared to cage odor (Fig. 6g). There was a significantly positive correlation between individual dendritic segment activity and spine stability (Fig. 6h; p = 0.009; Table 3), showing that pre-synaptic activity was sufficient to drive the stability of spines. This correlation, however, was transient as prolonged CNO exposure may have caused a network adaptation/homeostatic driven decrease in GC activity (Extended Data Fig. 9a), which explains the transient nature of the correlation between activity and stability (Extended Data Fig. 9c). A statistically significant correlation was seen on day 2 after CNO application, but was lost on day 3 (Extended Data Fig. 9d; day 1, p = 0.28; day 2, p = 0.009; day 3, p = 0.859). However, the correlation variability across days did not change after the persistent activation of MCs (Fig. 6i; *cage odor*: 0.396 ± 0.167; *CNO*: 0.348 ± 0.149, p = 0.352). Taken together, our data indicates that continuous pre-synaptic activation is sufficient to drive GC spine stabilization, suggesting that this component of the OB circuit primarily drives GC apical spine dynamics. These findings cannot rule out the role of feedback from the piriform cortex or other top-down inputs, therefore we pursued a computational model that was confined to the OB local circuit to determine the functional output.

### The activity-stability relationship, pattern separation, and persistent memory *in silico*

We developed a computational model to study OB network evolution resulting from spine structural plasticity. Since the *in vivo* measured correlation between spine stability and dendritic GC-activity was relatively low, but significantly positive, we tested whether such modest correlations can induce functionally relevant network connectivity and asked what function it may support. Our firing-rate model included excitatory MCs and inhibitory GCs interacting via reciprocal synapses (Fig.7a). The MCs received steady sensory stimulation (Fig. 7a, ‘Input Pattern’), leading to steady MC activity patterns that evolved slowly due to structural plasticity-driven reorganization.

**Figure 7:**
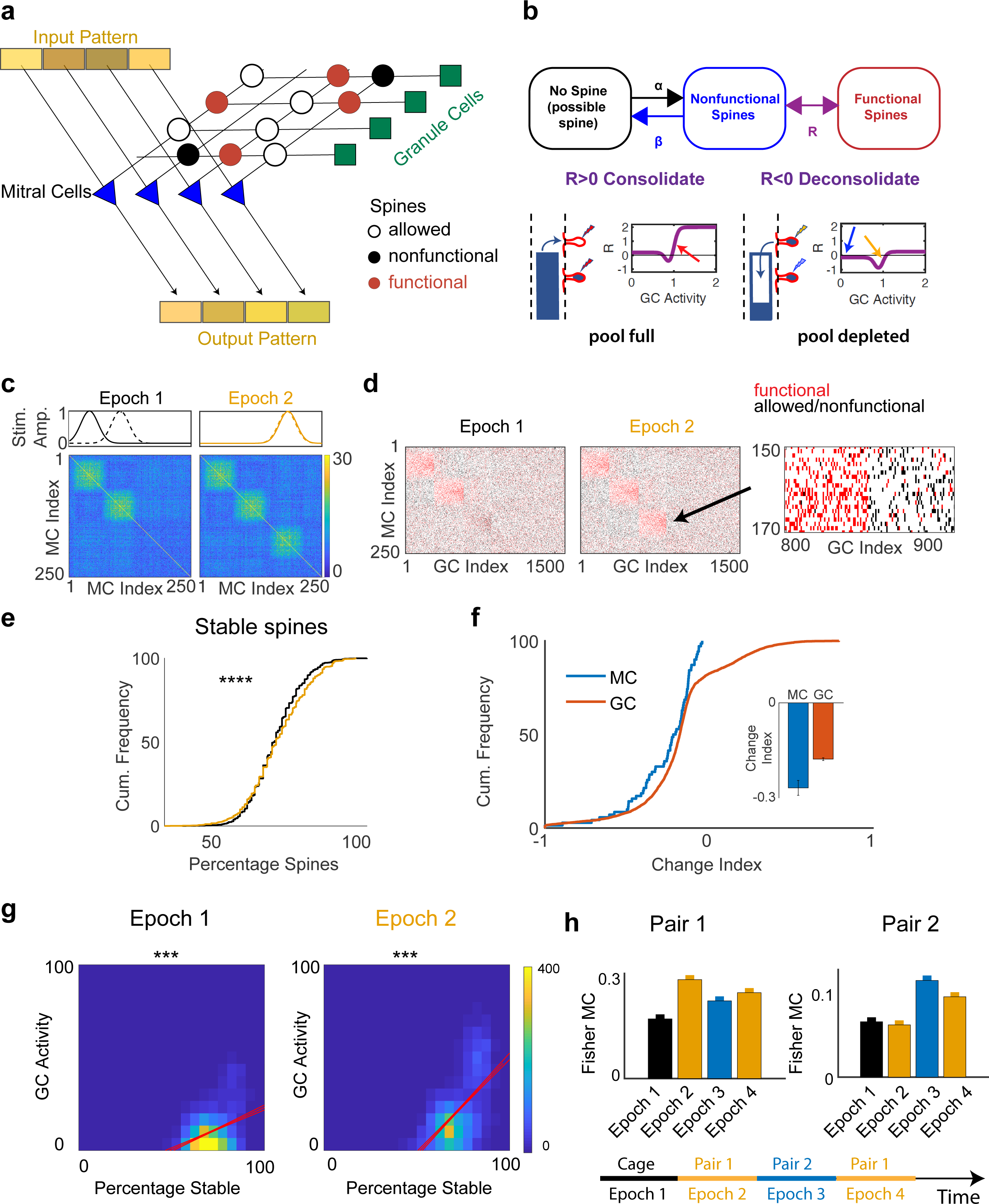
Computational model of spine dynamics. a. Sketch of the model network. Functional and nonfunctional spines are marked with red and black solid circles, respectively. Connections that are anatomically allowed but not realized, are marked with open circles. b. Two-stage model for spine formation. The transitions occur randomly with rates *α*, *β*, and *R*. Consolidation of spines requires large GC activity and a sufficiently filled resource pool (red arrow). Intermediate GC activity drives spine deconsolidation (orange arrow). c. Training with simple model stimuli induced disynaptic reciprocal inhibition of co-activated MCs the strength of which is given by the number of mediating GCs. d. Only functional MC-GC connections (red dots) mediated inhibition. e. Stimulus exposure increased the fraction of stable spines. The shifts in the cumulative distributions were small, but highly significant. f. Stimulus exposure reduced the MC activities more than those of GCs. g. Spine stability increased with GC activity, substantially more so after odor exposure in epoch 2 (cf. Fig.1i). h. Discriminability of the stimuli in pair 1 (left) and pair 2 (right). Pair 1 was used during epochs 2 and 4, pair 2 during epoch 3, as indicated on the timeline.

Our model is minimal and phenomenological in nature, motivated by various experimental observations ^18^. Our *in vivo* experiments showed a fraction of PSD-95-spines that are presumably nonfunctional (Extended Data Fig. 6). This was modeled with a two-stage process where initially nonfunctional (unconsolidated) spines were formed on a GC dendrite at allowed locations with a probability that increased with MC activity ^18, 26^. Subsequently, they were consolidated if the GC-activity was above a certain threshold (Fig. 7b).

We first illustrate the model using simple model stimuli (Fig. 7c). A statistically steady state was established through repeated exposure to a first stimulus set mimicking the experimental cage-odor baseline. Since the spine consolidation has Hebbian characteristics, connections were stabilized between coactive MCs and GCs (Fig. 7d). The number of such GCs was greatly enhanced between MCs that were co-activated by the presented odors (Fig. 7c bottom panels). In a second epoch, two very similar stimuli were used (same pattern with very slight offset; Fig. 7c) that drove an entirely different set of MCs. These new stimuli primarily activated a new set of GCs, which provided mutual inhibition mainly between MCs excited by the second stimulus pair (Fig.7c, d). Since the MCs that were stimulated during epoch 1 were not activated during epoch 2, the previously established connectivity was largely preserved.

Next, we used naturalistic stimuli based on glomerular activation patterns ^27^ (Extended Data Fig. 11a). A single stimulus in epoch 1 mimicked cage odor, followed by epoch 2 with two very similar stimuli to test whether network restructuring improves their discriminability. Overall, spines stabilized, decreasing the fraction of dynamic spines (Fig. 7e, Extended Data Fig. 14d). The cumulative distribution shift decreased with increasing dependence of the spine formation on MC activity (Extended Data Fig. 10). Comparison with the *in vivo* results suggested therefore that MC activity drives spine formation only to a limited degree.

The relative change in activity of each MC and GC over the course of epoch 2 (change index ^21^ was broadly distributed (Fig. 7f). Odors introduced in epoch 2 activated previously not activated MCs, resulting in new connections that activated numerous GCs (Fig. 7d), causing their change index to be positive. Overall, the mean of the change index was negative for MCs and less so for GCs, consistent with the *in vivo* results ^21^.

We then correlated the fractions of stable and dynamic spines of each GC with that GC’s activity (Fig. 7g, Extended Data Fig. 14a). Following epoch 2, the fraction of stable spines was positively correlated with activity (Fig. 7g), while the fraction of dynamic spines exhibited a negative correlation (Extended Data Fig. 14). During epoch 2 these correlations were more pronounced than in epoch 1, which was in part due to the increased stimulus intensity during the odor exposure.

To illustrate how the restructuring of the network reflected the training stimuli, MCs were sorted to separate the MCs that were predominantly excited by epoch 1 (‘cage odor’) from those of epoch 2 (Extended Data Fig. 14b). Since the odors in epoch 2 were added to the cage odor, significant connections were also established between the MCs excited by those odors and the MCs driven by the cage odor. Despite the substantial changes in the connectivity, the average number of consolidated spines per GC, as well as the fraction of consolidated spines, exhibited only a small, yet significant increase, which was consistent with our experiments (Extended Data Figs. 6, 13).

We used a Fisher discriminant to quantify the discriminability of very similar training odors. We found that the network restructuring significantly enhanced the discriminability of the odors driving the plasticity (Fig. 7h). Furthermore, we tested for “memory” in the learned network connectivity and the resulting odor discrimination ability. In an additional epoch 3 a new stimulus pair was presented (Extended Data Fig. 11b), while epoch 4 repeated the stimuli of epoch 2. As expected, epoch 2 led to a significant increase in the Fisher discriminant for the first, but not the second pair, while the Fisher discriminant for the second pair increased during epoch 3 (Fig. 7h). Importantly, during epoch 3, the Fisher discriminant for the first pair decreased only somewhat. Thus, despite the high dynamics of the structural plasticity that removed spines from moderately activated GCs (purple arrow in Extended Data Fig. 11c and Extended Data Fig. 12) and added spines to highly activated GCs (red and blue arrows), the network connectivity preserved the memory of the task involving the first stimulus pair, since epoch 3 did not activate the MCs involved in that task (yellow arrow).

## Discussion

GC dendritic spines are amongst the most dynamic in the brain and regional activity has been correlated with relative stabilization of these spines, yet the circuit mechanism driving this stabilization was unknown. For the first time, we show a direct correlation between the activity of individual GC dendritic segments, driven by persistent sensory stimulation, and the stability of their spines *in vivo,* over multiple days, and we provide evidence that this is directly driven by MC activity. Furthermore, our computational model shows this spine structural plasticity to be sufficient to increase discriminability while retaining sensory memory.

### Persistent odor exposure drives GC apical spine stability within active dendrites

We explored the dependence of an individual GC’s activity to drive its spine stability by combining structural and Ca^2+^ imaging *in vivo* in awake mice. Under baseline cage odor conditions, GC dendritic activity versus spine stability was uncorrelated, but increasing activity with chronic odor stimulation was sufficient to drive this correlation. This result is somewhat remarkable since spine stability was measured from ∼15-20 spines per segment, causing a ∼17 % increase in stable spines, indicating a change of ∼3 spines on the segment measured was sufficient for this correlated activity effect. These results support previous studies that showed *regional* OB activity was sufficient to drive spine stability ^17, 28^. Since spine stability has also been shown to correlate with learning ability in the somatosensory/motor cortex ^7, 8^, it suggests the activity-stability correlation we observed could be generalized to other neuron types from brain regions, which would be of interest for future studies.

To understand the mechanism of this activity-stability correlation, intracellular Ca^2+^ is important for spine formation and stability in cortical pyramidal neurons as shown in *ex-vivo* preparations ^29, 30^ where spine lifetime was positively correlated with Ca^2+^ peak frequency and duration. Chelating Ca^2+^ resulted in lower spine formation, spinule number and cumulative lifespan per spine ^29^. When we presented short duration odor, the activity-stability correlation was not induced, although this stimulus did shift the cumulative distributions of stable and new spines, but this did not correlate with individual dendritic activity. As to the cellular machinery of spine maintenance and formation, spine stability and lifetime are also associated with the gain of PSD-95 ^31^ and its expression complexity ^29^, respectively, whereas spine removal is often preceded by PSD-95 loss ^31, 32^. Although it has been shown that GC PSD-95 puncta stabilize with odor enrichment ^17^, suggesting an overall activity-dependence, our current results support a direct relationship between activity and stability.

We also explored how blocking sensory input affects the activity-stability correlation. Four days of naris occlusion greatly decreased GC activity, but, compared to the cage-odor condition, spine stability was not significantly changed, while there were fewer new spines and more lost spines. Thus, during deprivation, it is possible that some of the strong GC synaptic connections formed during continuous odor exposure may remain stable, with evident pruning in parallel. This may be a hallmark of inhibitory neurons since in the visual cortex monocular deprivation also caused increased inhibitory synapse loss and decreased synapse addition in the dendritic shafts of layer 2/3 pyramidal neurons, whereas layer 2/3 pyramidal neuron spines were not affected ^33^, but a significant increase in spine formation occurred in layer 5 pyramidal neurons ^34^. Therefore, spine changes appear to be neuron-type specific and may reflect network configuration and input differences across brain regions.

### Spine activity-stability relationship depends on dendritic location

We also measured the spine dynamics of GC proximal spines which receive unidirectional top-down cortical input. In contrast to the distal GC spines, which primarily make reciprocal dendro-dendritic connections with MC cells, the proximal spines did not have an activity-spine stability relationship with continuous odor exposure. This supports region-specific spine adaptability to differential types of input, as shown in other studies: For distal spines, spine density increased with simple olfactory enrichment ^35, 36^ and PSD-95 spine density decreased with deprivation, whereas proximal spine density increased ^37^. Also, a more complex input, by pairing an odorant with a reward task, caused increased proximal spine density with no effect on distal spines ^38^. Associative learning induced an increase in GC excitatory responses, increased spine formation and stabilization, and improved odor discrimination. In parallel, an associative discrimination tasks also increase the cortical feedback to the GCs, and was suggested to provide contextual information to drive GC context dependent plasticity ^9^. Therefore, these compartment-specific features may provide an additional level of segregation of synaptic inputs and therefore increase the adaptability of GCs.

### MC activation is sufficient to drive activity-stability relationship

We dissected the OB circuit to find whether intrinsic components are sufficient to drive the activity-stability relationship of GC distal spines. Cell-autonomous silencing of GCs was performed and no activity-stability relationship was found. This finding is supported by other studies showing network silencing having no effect on hippocampal excitatory and inhibitory synapse density ^39^, spine volume fluctuations in small spines ^40^ or change in gephyrin inhibitory puncta dynamics ^41^. Additionally, we performed cell-autonomous GC excitation which did not cause an activity-stability relationship in the distal spines. This is supported by histological analysis where GC excitatory synapses across all compartments were shown to have no change in density with intrinsic excitation ^37^. Since increased sensory input was the only condition that drove the activity-stability relationship, this suggested that pre-synaptic activity of MC-cells may provide a critical role.

Previously we demonstrated that gephyrin puncta dynamics on MC cell lateral dendrites match GC spine dynamics ^10^. We proposed a Hebbian plasticity mechanism to stabilize odor representations and facilitate pattern decorrelation between GC-MC cell synapses. In the current study we performed chronic MC-cell autonomous activation to determine if this was sufficient to drive the activity-spine stability relationship. MC-cell activation was able to drive GC activity-dependent spine stability, similar to that of sensory stimulation, albeit transiently. This is in agreement with a study showing MC-cell stimulations, mimicking odor activation, stabilized dynamic filopodia-like structures on spines by increasing their lifetime and reducing their mobility and was shown to be dependent on MC glutamate release and BDNF ^26^. This effect was observed in 45-minute sessions with glutamate stimulation, whereas we observed spine stability persistence over 3-days with MC activation, suggesting a conserved mechanism. However, we observed the GC activity-spine stability relationship in the initial days of DREADD-activation of MC cells, but the relationship was progressively lost. Further study with more temporal control methods for disrupting specific pre- and post-synaptic components *in vivo* would be interesting to determine what patterns of activity are sufficient to mimic the sensory input driving GC spine stability.

### Minimal computational model of spine plasticity in the MC-GC circuit: odor discrimination memory

We utilized a minimal computational model to provide insight into the *in vivo* results for spine plasticity. *In vivo* and *in silico* continuous odor presentation increased spine stability that positively correlated with the activity of the dendrite associated with the spine. Odor exposure led only to a small increase in the fraction of consolidated spines, which was consistent with our *in vivo* result showing a small decrease in PSD^+^ -spine fraction resulting from odor deprivation. Nevertheless, in the model this small increase indicated a substantial restructuring of the network.

Due to the network reorganization caused by odor exposure, the MC and GC activities evolved in the model. On average, their activity decreased with odor exposure ^21^. However, while the activity of almost all MCs decreased, a significant number of GCs were newly recruited implying an increase in their activity. The network restructuring led to the formation of new subnetworks in which GCs provided disynaptic inhibition among MCs that were strongly activated by the odor during odor exposure. This specific inhibition enhanced the discriminability of very similar stimuli related to that odor ^42^, consistent with previous experiments showing perceptual learning in a habituation task ^43^ that required the activation of partially overlapping regions in the OB ^43^. This was also the case in a computational model implementing synaptic-weight plasticity ^43^.

An important aspect of our model is its memory for learned odors ^44^. In sequential exposure with multiple odors, previously learned connectivities were only compromised if the new stimuli significantly overlapped with the previous ones. Then, the bulbar odor representations of the previously learned stimuli were modified, resulting in a partial loss of the memory of the previously learned discrimination task. For a small overlap, however, odor memories persisted. For the stimulus mimicking “cage odor”, the model exhibited a significant correlation, albeit weaker than for the odor-exposure stimulus, while experimentally the correlation was not significant. This apparent discrepancy may be due to mice experiencing the same cage odor throughout their lives.

In our *in vivo* experiments, a large fraction of dendrites with high activity did not form new spines (Fig.1i). The model showed that this does not imply any direct influence of GC activity on spine formation; instead, it was sufficient if high-activity GCs had a large number of consolidated spines, *relative* to which the number of new spines was small (Extended Data Fig. 12a bottom panels), therefore the consolidation of the spines depended crucially on GC activity.

A prominent feature of the OB circuit is the extensive top-down projections that predominantly target GCs, allowing higher brain areas to modify OB odor processing. Previous modeling suggests that GC adult neurogenesis naturally leads to a network structure enabling higher brain areas to inhibit specific MCs that allows context-dependent odor-processing in the bulb ^45^. Since the GC spine plasticity mimics neurogenesis by sharing the an almost unlimited level of plasticity, albeit local activity/plasticity versus an entirely newly integrated cell, it is expected that mature GC spines contribute by providing specific control of bulbar processing influenced by higher brain areas.

Our studies demonstrate a potential mechanism for perceptual learning during continuous odor exposure based on activity-dependent structural plasticity of adult-born GCs. While recent studies highlight the strong enhancement of structural spine plasticity by cortical contextual feedback during active learning, we show that presynaptic activity is necessary and sufficient to drive the correlation between individual GC activity and spine stability.

Therefore, this enables the formation of stimulus-specific subnetworks within the OB to enhance stimulus discriminability, highlighting a perceptual learning role that the local OB circuit provides.

## Acknowledgements

We thank Matt Valley for his contribution in developing root components of the calcium imaging code, Gabriel Lepousez and Sebastian Wagner for assisting in the development of the awake imaging setup and odor delivery system. Timothy Sailor for machining essential parts of the mouse imaging stage, Anne Lanjuin for providing Tbet-Cre mice, Adi Mizrahi for providing the PSD-95-GFP construct, Florian Ruckerl and David DiGregorio for managing the Pasteur Institute Neuroscience Department imaging facility. We thank Sanjana Sebastian and Jorge Soriano Campos for assistance with experiments. Soham Saha’s PhD fellowship was funded by the École Neurosciences de Paris (ENP) network. This work is supported by the life insurance company “AG2R-La-Mondiale“, Agence Nationale de la Recherche (ANR-15-CE37-0004-01), Agence Nationale de la Recherche (ANR-15-NEUC-0004, Circuit-OPL) and the Laboratory for Excellence Programme “Revive”. Also funded by the National Institutes of Health, www.nidcd.nih.gov, under grant R01-DC015137 (to HR).

## Author contributions

Conceptualization, K.A.S, S.S, H.R. and P.M.L.; Methodology, K.A.S., S.S, J.H.M., H.R. (computational model); Investigation, S.S., G.A., J.H.M. (computational model); Formal analysis, J.H.M. and H.R. (computational model); Formal analysis, S.S., G.A.; Writing – original draft, S.S., J.H.M., H.R. and K.A.S.; Writing – review and editing, K.A.S., P.M.L. and H.R; Supervision, K.A.S, P.M.L. and H.R.

## Methods

### Animals

Eight-week old C57Bl/6j (Janvier Laboratories, Le Genest-Saint-Isle, France) and Tbet-cre mice (C57Bl/6j background, 8 weeks old) ^25^ housed under standard conditions were used in the study. All experiments were performed in compliance with the French application of the European Communities Council Directive (2010/63/EEC) and approved by the local animal welfare committee of the Institut Pasteur (CETEA, project #2013-030).

### Stereotaxic injections

OB granule cells (GCs) were labelled at a specific age by injecting lentivirus expressing a Cre-recombinase with a tdTom reporter under the CMV promoter (LV-CMV-tdTom-IRES-Cre) ^46^ into the rostral migratory stream (RMS) as described previously ^10^. Mice were anesthetized (150 mg/kg ketamine, 5mg/kg xylazine; 0.02 mg/kg buprenorphine) and the head was secured to a stereotaxic frame (David Kopf) while the body temperature was maintained using a rectal temperature feedback heating pad. Using sterile technique, the skull was exposed and 0.5 mm craniotomies were drilled for bilaterally injecting viral solution into the RMS using a glass micropipette attached to a Nanoinjector system (Drummond Scientific) at the following coordinates relative to the bregma: 3.3 mm anterior, ±0.82 mm lateral, and 2.9 mm deep. The scalp was sutured and the mice were returned to their home cages with free access to food and water.

After 3 weeks, once the Cre-labelled neuroblasts migrated to the OB, floxed adeno-associated virus (AAV) expressing the Ca^2+^ indicator GCaMP6f was injected in the OB (AAV-hSyn-DIO-GCaMP6f-WPRE), either alone or in combination with a floxed AAV expressing excitatory or inhibitory DREADD (Designer Drugs Exclusively Activated by Designer Drugs, AAV5-hSyn-DIO-hM3D(Gq)-mCherry-excitatory; AAV5-hSyn-DIO-hM4D(Gi)-mCherry-inhibitory). This labelling strategy permits either recording GC activity (with GCaMP6f) or to activate/suppress GCs in a temporal manner by administering the ligand for DREADD, Clozapine-N-oxide: CNO, thereby having specific action on neurons co-expressing Cre and DREADD. To label GC complete dendritic/spine structure and post-synaptic excitatory synapses, a lentivirus expressing tdTom (LV-hSyn-tdTom) and a lentivirus expressing a post-synaptic density-95 (PSD-95) GFP fusion (PSD-LV-CMV-PSD-95-GFP (Mizrahi, 2007), respectively, were injected using the same coordinates as described above. Transgenic mice with the expression of Cre under the Tbx21 promoter (Tbet-Cre) (Haddad et al., 2013) were injected with floxed GCaMP6f (AAV-hSyn-DIO-GCaMP6f-WPRE) at the same coordinates as indicated above, causing expression to be restricted to mitral/tufted cells (MCs) for recording their neuronal activity. The details of the quantity of viruses injected, brain region infected and viral titre are summarized in table 1.

**Table 1:** List of viruses used for labeling neurons.

| **Viral construct** | **Purpose** | **Viral titre** | **Region** | **Cat. #** | **Vol. (nl)** | **Supplier** |
| --- | --- | --- | --- | --- | --- | --- |
| LV-CMV-tdTomato-Cre | Labeling adult born GCs at a particular age, structural marker | $6 \times 10^8$ VP/ul | RMS | | 200 | ICM Vector Platform, Paris France |
| LV-CMV-GCamp6f | Co-injected with LV-tdTomato-cre to label adult born GCs, $\text{Ca}^{2+}$ sensor as a neuronal activity marker | $1.26 \times 10^9$ VP/ul | RMS | SL100322 | 250 | SignaGen |
| LV-EF1a-PSD95-GFP | Label adult born GC glutamatergic synapses as a proxy of functional spine turnover | $1.38 \times 10^8$ VP/ul | RMS | | 200 | ICM Vector Platform, Paris France |
| AAV9 Syn-FLEX-GCamp6F.WPRE.SV40 | Synapsin-driven, Cre-dependent $\text{Ca}^{2+}$ sensor; activity marker for cre-labelled adult born GCs | $7 \times 10^{13}$ VP/ul | OB | 100833 | 500 | Addgene |
| AAV5-hSyn-DIO-hM3D(Gq)-mCherry | DREADD receptor with mCherry reporter for CNO-induced cre-labelled adult-born GC activation | $7 \times 10^{12}$ VP/ul | OB | 44361 | 300 | Addgene |
| AAV5-hSyn-DIO-hM4D(Gi)-mCherry | DREADD receptor with mCherry reporter for CNO-induced cre-labelled adult-born GC suppression | $4.5 \times 10^{12}$ VP/ul | OB | 44362 | 300 | Addgene |
| AAV8-CaMKIIa-hM4D-mCherry | DREADD receptor with mCherry reporter for CNO-induced GC suppression | $4.41 \times 10^{12}$ VP/ul | Olfactory cortex | | 300 | Univ North Carolina, USA |

### Cranial window procedure

At the timepoint for OB floxed virus injections, a cranial window was implanted for chronic *in vivo* imaging. A craniotomy was performed by carefully cutting the skull with a #12 scalpel blade slightly larger than the cover glass dimension (3.0 × 1.4 mm) over the OB. The skull was removed and a custom cut glass cranial window (UW-Madison microelectronics from my paper) was placed in the craniotomy and the glass-skull interface was sealed with dental cement (Metabond). A custom-made stainless-steel/brass head bar (0.40g) was secured to the skull with cyanoacrylate glue and the exposed skull surface was covered with dental cement. The mice were house individually and placed on antibiotic-water (Avemix, 8g in 1000ml water) for 5 days after the cranial window surgery. For the duration of the imaging experiments the animals had free access to food and water.

### *In vivo* two-photon imaging

#### Plane of imaging

The somas of GCs in the OB are broadly distributed in the granule cell layer (GCL; ∼500 µm thick layer) with their apical dendrites projecting to the external plexiform layer (EPL) where they make dendrodendritic synapses with the MC cells ^11, 47^. GCs were sparsely labelled as required for accurately tracing and measuring GC structural changes. With sparse labelling and random insertion of GCs in the thick GCL, it was necessary to perform calcium imaging on GC dendrites. This provided three advantages: 1) A large number of GCs can be sampled for calcium imaging in one z-plane which was not possible in the GCL with sparse labeling, There is no bias in imaging GCs confined to specific GCL planes and 3) Individual GC branch activity can be directly correlated with its individual spine dynamics. The EPL was imaged to capture structural and activity changes in GC apical dendrites and the internal plexiform layer (IPL)-GCL for imaging GC proximal dendrites (**Extended Data** **Fig. 3a, Extended Data** **Fig. 4a-b**). Only dendrites which were horizontal to the imaging plane were used for structural tracing to avoid z-spread artefacts. For recording neuronal activity in the MCs, mitral cell somas were imaged in the mitral cell layer (MCL).

#### Awake 2-photon imaging

Calcium activity in the GC dendrites were imaged using a two-photon Prairie Investigator microscope (Bruker) with a resonant galvanometer attached to a DeepSee Ti:Sapphire femtosecond pulsed laser (Mai-Tai, Spectra Physics) to excite GCaMP6f at 950 nm. Images were acquired using a 20X 1.05 NA objective (Olympus). The same region was found between days using anatomical landmarks including blood vessel and dendritic projection patterns for reference. Time series were recorded at 512 x 512 pixels at 15 Hz. Calcium activity was either recorded spontaneously (2000 frames per trial) or in odor evoked condition (300 frames per trial).

#### Odorant delivery

Iso-amyl acetate (IAA, Sigma Aldrich, Germany) odorant was used as the monomolecular odorant for odor exposure which was diluted 1:10 in mineral oil for presenting to the mice. For odor evoked calcium recordings, IAA was further diluted with humidified air (1:10) and presented to the mice directly in a mask surrounding the nose for 2s per trial using a custom-built olfactometer that was adapted for head-fixed imaging (Alonso et al., 2012). Odors were delivered with 2-3 blocks of 20 trials were recorded per day per animal.

For odor exposure timepoints, IAA was absorbed on paper in tea balls and placed in the animal cage above the cage grid, as described previously (Moreno, PNAS, 2009; Mizrahi, Nat Neuro, 2011) and the odorant was present for 24 h during 4 days.

#### Naris occlusion protocol

The construction of nose plugs for unilateral naris occlusion was followed based on the method described in Cummings et al., 1997 ^48^. Polyethylene (PE-10) tubing (Becton Dickinson, Parsippany, NJ), a silk surgical suture, and human hair. The human hair was tied to an end of the suture and then threaded into the PE tubing. The suture was passed into a knot around the tied end of the hair and passed into the lumen of the tubing. Finally, after the knot was slid into the lumen, the tubing and the suture was trimmed into roughly 0.5cm, ensuring that a small part of the hair remained outside so as to allow us to remove the plugs after experimentation. One end of the plug was trimmed to make it into a sharp end to make it easier for the nose plugs to be inserted into the nose of the mice. Mice were anesthetized temporarily using isofluorane, and the plugs were placed in the nose as and when described during the course of the experiment. Animals were unable to remove the nose plugs during the time of the experiment. They were returned to their home cages with ready access to food and water, and were under constant supervision during the whole course of the deprivation study.

### DREADD activation and inhibition

Virus expression of DREADD, as described above, was used for selective continuous activation or suppression of neuronal activity ^49, 50^. Mice either were injected with excitatory or inhibitory DREADD viruses exclusively in the GCs or in the MCs. Clozapine-N-oxide (CNO, 0.025 mg/ml in water, selective for hM4Di and hM3Dq; Sigma-Aldrich) was administered in drinking water based on daily water intake of 6 ml, providing 5 mg/kg/day of CNO.

### Neuronal activity analysis

#### Identifying z-plane of imaging between sessions

A semi-automatic method was developed to consistently find the same imaging z-plane between imaging sessions. The imaging z-plane was determined on the first day of imaging and maintained throughout all session using the tdTom channel as a structural reference (Extended Data Fig. 2a-b). For subsequent imaging sessions, the animal was positioned in the same fixed location on the imaging stage and landmarks (blood vessels, dendritic structure etc.) were used to visually estimate the target z-plane. A 31 µm z-stack was acquired, ±15 µm above and below the estimated z-plane (Extended Data Fig. 2b). Using custom code developed in Matlab (Mathworks) the first session target z-plane was normalized and registered to the subsequent imaging session estimated z-plane (StackReg, ImageJ). A 2-dimensional cross-correlation was performed between each z-plane of the subsequent imaging session z-stack and the target normalized/registered z-plane, as the reference (Extended Data Fig. 2b). The z-plane with the highest correlation was used for the imaging session.

#### Animal movement filtering and image registration

The acquired images were grouped into blocks of odor evoked trials with each block consisting of 20 trials, each containing 300 frames (22s) and spontaneous activity was grouped into blocks of 2, each containing 2000 frames (132s). Since the imaging of activity was performed in awake mice, it was necessary to correct for lateral and out-of-frame z

movements. For each imaging block, lateral x-y-movements were corrected on the tdTom channel using rigid registration (MultiStackReg, ImageJ, [http://bradbusse.net/sciencedownloads.html]) and the registration transformation was applied to the GCaMP6f channel.

For the elimination of frames with z-displacements out of the target z-plane, within each block, a mean image was calculated for all the tdTom frames which was used as the target z-plane reference. A 2-dimensional cross-correlation was performed on each tdTom image in the time series versus the target z-plane reference (Matlab, Mathworks; Extended Data Fig. 2c). A manually defined threshold was defined at the lower tail of the histogram distribution of the correlation coefficients (Extended Data Fig. 2d). The frames identified with out-of-frame movements on the tdTom channel were indexed and removed from the GCaMP6f channel and replaced as NaN values Extended Data Fig. 2e. The entire trial was removed if the percentage of dropped frames exceeded 25% of the total frames.

Each block was also x-y-registered to correct for between session/day drift. The average tdTom T-projection of each block was affine registered to the first imaging session (MultiStackReg, ImageJ; Extended Data Fig. 2f-g) and the transformation of the block was applied to each image within the block of the GCaMP6f channel.

#### Region of interest detection

For increasing the signal-to-noise ratio and diminish principal component (PC) filtering was performed using the first 25 principal components. PCs were assigned from the raw images based on how pixels in the imaging field covary in the time dimension and with an orthogonal transformation, the observations were converted into linearly uncorrelated variables. This method effectively removed static background and filtered noise since the non-varying pixels and low amplitude random signal, respectively, have low variability across time, thus the image was reconstructed using the highest variable eigen vectors (Extended Data Fig. 2H, I). The filtering used the *pca* function in Matlab from the Signal processing toolbox (Mathworks Inc.).

Elliptical ROIs were manually traced in FIJI (ImageJ) on the dendritic fragments of the PCA-reconstructed images (Extended Data Fig. 2I, J). The PCA-reconstructed images were concatenated across multiple blocks to aid in visualizing segments that flashed (Extended Data Fig. 2J). In parallel, a standard deviation projection was used to highlight the pixels with the highest variance. The ROI coordinates were imported with custom made code in MATLAB using the Miji interface and ReadImageJROI code (https://www.mathworks.com/matlabcentral/fileexchange/32479-readimagejroi). For minimizing neuropil contamination and noise from each ROI, PC filtering was applied to the pixels within each individual ROI (Extended Data Fig. 2J), using the linear combination of reconstruction as previously described, and pixels within the first PC were used to define the ROI boundary (Extended Data Fig. 2J). This filtering increased the robustness for activity quantification by eliminating pixels containing noise thereby increasing the signal-to-noise ratio by ∼25% (Extended Data Fig. 2K-M). For ROIs below x threshold, thereby having no activity for a given block, the original elliptical ROI was used. The PC filtered ROI coordinates were used to extract the mean area intensity for each frame indexed dropped frames, as previously determined, were filled with NaN values.

#### Calcium activity normalization and significance test

For each ROI, the pixel intensity values across time were smoothed across 5 frames using the *NaNfastsmooth* function in MATLAB. The fluorescent signal was normalized using the ΔF/F ratio:

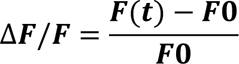

*F0* is the average pixel intensity over the entire trial period or the mean of the pixel intensity values before odor delivery for spontaneous and odor evoked trials, respectively, with *F(t)* being the fluorescent intensity (*F)* at time *t*. Significance of evoked activity for each ROI was determined by the Wilcoxon ranksum test comparing between the pre-evoked (1 to 7 s) and evoked (8 to 10 s) frames for each trial, selecting trails where p < 0.01 was considered a significant response.

*Area under the curve (AUC) calculations*:

The magnitude of GC responses of for each ROI are represented by the area under the curve (AUC) of the ΔF/F after the initiation of the odor presentation and in the entire trial for odor evoked and spontaneous trials, respectively. The *trapz* and *nansum* MATLAB functions were used to calculate the AUC values. A non-parametric KS test was used to compare the differences in the cumulative distribution with a p-value of less than 0.05 considered significant.

### *In vivo* two-photon imaging protocol of GC dendrite structure

#### Anaesthetized imaging of GC dendrites

The first imaging session was at least 4 weeks after the cranial window surgery to insure inflammation from the procedure had subsided ^10^. Anesthetized imaging was performed for the co-localization studies of PSD-95 and tdTom in dendritic spines and for short-term dynamics to compare with awake-state dynamics. Isofluorane was used for anesthesia (0.8% in oxygen) and body temperature was maintained at 37° C with a rectal thermometer feedback-heating pad. The same 2-photon setup was used, as indicated above, but for PSD-95 GFP and Td tomato co-localization a 2X digital zoom was used for 0.22 x 0.22 x 2 µm (x,y,z) resolution stacks.

#### Awake imaging of GC dendrites

For *in vivo* imaging of GC dendrites, the same imaging setup was used. Ten Serial stacks (0.44 x 0.44 x 2 µm [x,y,z]) of the same volume were rapidly acquired using the resonant galvanometer scanner (period 0.06595s per slice, taking into account the z-step motor). This oversampling of the same volume allowed for filtering out imaging planes with animal movement artefact (Extended Data Fig. 1).

### Structural reconstruction of dendrites in awake-imaged volumes

A protocol was established to reconstruct awake-imaged volumes to have sufficient resolution to track dendritic spines. The main steps of this protocol are summarized in Extended Data Figure 1. Structural imaging in awake animals comes with two major dimensions of movement: translational x-y movements and out-of-plane z movements (Extended Data Fig. 1a). While translational full frame x-y shifts can be corrected using available registration algorithms ^51^, out-of-plane z movements need to be discarded due to combined loss of information from the target plane and oblique warping in the z-axis (Extended Data Fig. 1b). Since mouse movement was relatively sparse and random, serial stacks were acquired to minimize movement artifact being concentrated on a particular slice within the volume. For the 10 serial acquired stacks, each frame was acquired in ∼66 ms, thereby limiting within-frame movement artifacts. Two-dimensional cross-correlation of the frames was used to preserve frames without movement and discard those with artifact (Extended Data Fig. 1b). The mean of the replicate frames that were without artifact were used to reconstruct the final volume. This protocol used the following detailed steps:

1. An arbitrary assigned series of stack reference x-y-slices were defined at a 10-15 frame interval (Extended Data Fig. 1b-d).
2. A 2-D cross-correlation (x-y) was calculated for every x-y-slice in the volume versus each of the reference x-y-slices (*corr2* function, MATLAB; Extended Data Fig. 1d).
3. The correlation values were smoothed (*smooth* function with 5 frames, MATLAB) and subtracted from an ideal Pearson’s correlation coefficient normal distribution.
4. For z-slices with subtracted correlation values over a median threshold, indicating significant movement artifact, the x-y-slice was indexed and replaced with a nan value, thereby excluded from the final reconstruction (Extended Data Fig. 1c, e).
5. The mean of each x-y-slice that was below the subtracted correlation threshold, for each corresponding volume, was used to reconstruct the final volume (Extended Data Fig. 1e-f).

This method allowed reconstructing the GC structure with high fidelity to allow tracking the same spine population across multiple days (Extended Data Fig. 1h). The mean percentage of dropped frames was 6.0% ± 0.6% and was insignificant across trials (Extended Data Fig. 1g; ANOVA across trials, F(9, 990)= 1.5; p = 0.242; adjusted p = 0.142). This reconstruction allowed tracking dendrites across multiple days in awake mice with high resolution and only slight variations in noise across imaging planes (Extended Data Fig. 1h). The percentage of dropped z-slices was ∼5 percent across animals (Extended Data Fig. 1d, f, g).

### GC spine tracking

The dendritic structures of the GCs were traced in 3-D using semi-automatic filament tracing (Imaris, Bitplane) while visualizing the structure with 3D glasses (3D vision, NVIDIA) as previously published (Sailor et al., 2016). Dendritic spines were termed “spines” in a broad definition, which includes a range of morphologies from mushroom to filopodia-like spines. Due to z-spreading issues with 2-photon imaging, only non-overlapping dendritic segments that were parallel with the imaging plane were traced and spines that projected into the z-axis were not traced. To aid visualization, a “fire” heat map was used to expand the dynamic range of the images. Images were traced with the previous day’s image adjacent for confirming if each spine was significant. PSD-95 puncta were marked with “spots” on the tdTom traced spines. Projected 2D images of the tracings were made, registered (StackReg, Fiji), arrayed in Illustrator (Adobe) 2D and horizontal lines were drawn to track spines across all timepoints (fig. sX). The tracked images were then automatically analyzed using custom Matlab code producing a table of stable, new and lost spines at each time-point.

The percent total spines were calculated as stable: 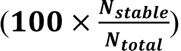, new: 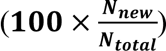 and lost: 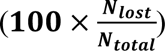, where Ntotal is the total number of spines at a given time (Nnew+Nstable+Nlost). Data are presented as mean ±95% C.I. and statistical analyses across spines of different days were performed using one-way ANOVA with Prism (Graphpad Prism). For distribution estimates, cumulative frequency (as percentages) representation was used along with the KS test to determine normality. The Pearson’s correlation coefficient was used as a measure of similarity between observed parameters across the experimental set-ups. A 5% confidence interval was considered for all analysis and p < 0.05 was taken as statistically significant.

### Correlating spine turnover and activity

#### Identifying dendrite-ROI pair

In order to compare how activity in a dendritic segment impacted the spine turnover across the particular dendrite, the ROI map for activity was overlaid at the proper z-level in the imaged structure volume in Imaris as a separate channel. Each particular traced dendritic structure was then manually assigned to a specific ROI for all the ROIs in the plane.

#### Spectral separation for DREADD activation/suppression

The excitatory and inhibitory DREADD AAV contains a mCherry reporter and GC structure was labeled with tdTom. To separate these red fluorescent signals in animals that had this labeling combination, z-stacks were acquired at 950 nm which excited both tdTom and mCherry and also acquired at 1020 nm which only excited mCherry. The pixel intensities of the two stacks were normalized and the 1020 nm acquired stack was subtracted from the 950 nm stack to expose the tdTom signal (Extended Data Fig. 8a-b). Using these combined stacks, tdTom^+^/GCaMP6f^+^ and tdTom^+^/GCaMP6f^+^/mCherry^+^ signals were discriminated to indicate which cells express DREADD. This resulted in an attrition of data by 2-4% of the original dendrite tracing (Table 2).

**Table 2:** Statistical power estimation for different experiments.

| Experiment | Statistical test performed | Number of animals used | Dendrite-ROI pairs tracked | Confidence Interval | Effect size (d) | Standard deviation | Difference in mean | Slope of curve | Attrition rate | Achieved Power |
| --- | --- | --- | --- | --- | --- | --- | --- | --- | --- | --- |
| Neuronal activity in odor exposure | K-S test | 8 | 51 | 0.05 | 0.72 | 0.19 | 0.135 | - | 0 | **0.95** |
| Spine stability in odor exposure | K-S test | 8 | 51 | 0.05 | 0.38 | 0.165 | 0.06 | - | 0 | 0.76 |
| Neuronal activity-spine stability correlation: distal dendrites | Linear regression; variation of slope of linear fit | 8 | 51 | 0.05 | 0.13 | 0.20 | 0.015 | 0.22 | 0 | **0.93** |
| Neuronal activity-spine stability correlation: proximal dendrites | Linear regression; variation of slope of linear fit | 4 | 55 | 0.05 | 0.06 | 0.17 | 0.24 | -0.03 | 0 | **0.81** |
| Neuronal activity-spine stability correlation: passive odor exposure | Linear regression; variation of slope of linear fit | 5 | 42 | 0.05 | 0.15 | 0.16 | 0.23 | 0.012 | 0 | 0.74 |
| Neuronal activity-spine stability correlation: Inhibitory DREADD | Linear regression; variation of slope of linear fit | 5 | 49 | 0.05 | 0.24 | 0.173 | 0.032 | 0.03 | 12% | **0.83** |
| Neuronal activity-spine stability correlation: Excitatory DREADD | Linear regression; variation of slope of linear fit | 4 | 36 | 0.05 | 0.01 | 0.153 | 0.004 | -0.23 | 16% | 0.75 |
| Neuronal activity-spine stability correlation: Naris occlusion | Linear regression; variation of slope of linear fit | 7 | 53 | 0.05 | 0.05 | 0.22 | 0.13 | -0.14 | 36% | 0.68 |
| PSD-95 turnover | K-S test | 5 | 33 | 0.05 | 1.25 | 0.22 | 0.13 | - | 24% | **0.82** |
| Neuronal activity-spine stability correlation: MC excitation | Linear regression; variation of slope of linear fit | 4 | 23 | 0.05 | 0.294 | 0.18 | 0.18 | - | 0 | **0.95** |

#### Min-max normalization of the AUC values

Neuronal activity is represented as under the curve (AUC) of significant calcium transients. Raw AUC values extracted from the time series were normalized by min-max normalization to scale the data from 0-1. The following formula was applied:

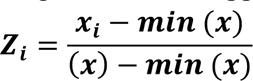

where, *Z* is the normalized value of the AUC raw value of *x* and *i* indicates the index of the dendrite-ROI pair. The maximum value in each animal was taken for the normalization.

#### Change index

To normalized the change in the parameters being recorded (neuronal activity or spine dynamics), the magnitude of change in the neuronal activity and percentage stability of the spines in a given dendrite-ROI pair was determined, giving the “change index” as summarized below:

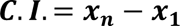

where, *C.I.* is the change index, *x* is the parameter whose change was quantified (min-max normalized neuronal activity or the percentage stability of spines) and *n* is the last day of the experimental condition (i.e., continuous odor exposure, passive odor exposure or chemogenetic manipulations).

#### Linear regression

As a means to represent coincidental change in neuronal activity and spine stability, linear regression was performed on both parameters. For each experimental condition, the data for these parameters were pooled across experimental days and represented per condition.

#### Euclidean distance

In order to account for the variability of the dendrite-ROI pairs in the activity versus stability scatter plot from the mean activity-mean stability, we used the Euclidean distance method, in Cartesian coordinates, if (x2, y2) and (x1, y1) are two points in two-dimensional Euclidean space, the distance between the two points is given by the formula: 

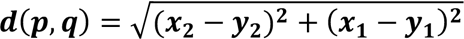

### Statistical analysis

#### Test for comparison of groups

Comparison of the data across groups was primarily done on the distribution of the data using the KS test. The non-parametric Wilcoxon Signed rank test was used for comparing the mean of neuronal activity or spine dynamics across two groups. For comparisons across days, the mean rank of each data per day was compared to the mean rank of every other day. For comparison across multiple groups, a non-parametric ANOVA test was used, followed by post-hoc Dunn’s correction. For the determination of the normality of the data, a Shapiro-Wilk test was performed. For comparing the regression fit, the variation of the slopes of the fit were compared to the internal control for each experiment. A p-value of 0.05 was considered significant.

#### Regression and correlation

As discussed previously, linear regression between activity in a small part of the traced dendrite was compared to the percentage of stable and dynamic spines of the entire dendrite. The p-value represents the validity of the null hypothesis that the existing slope of the regression varies significantly from 0. We have reported the r^2^ values for each of the regression plots, with the p-value computed from the slope of the fit. The Spearmans’ r and regression slopes for each experiment are reported in Table 3. Values in bold indicate significant observations.

**Table 3:**
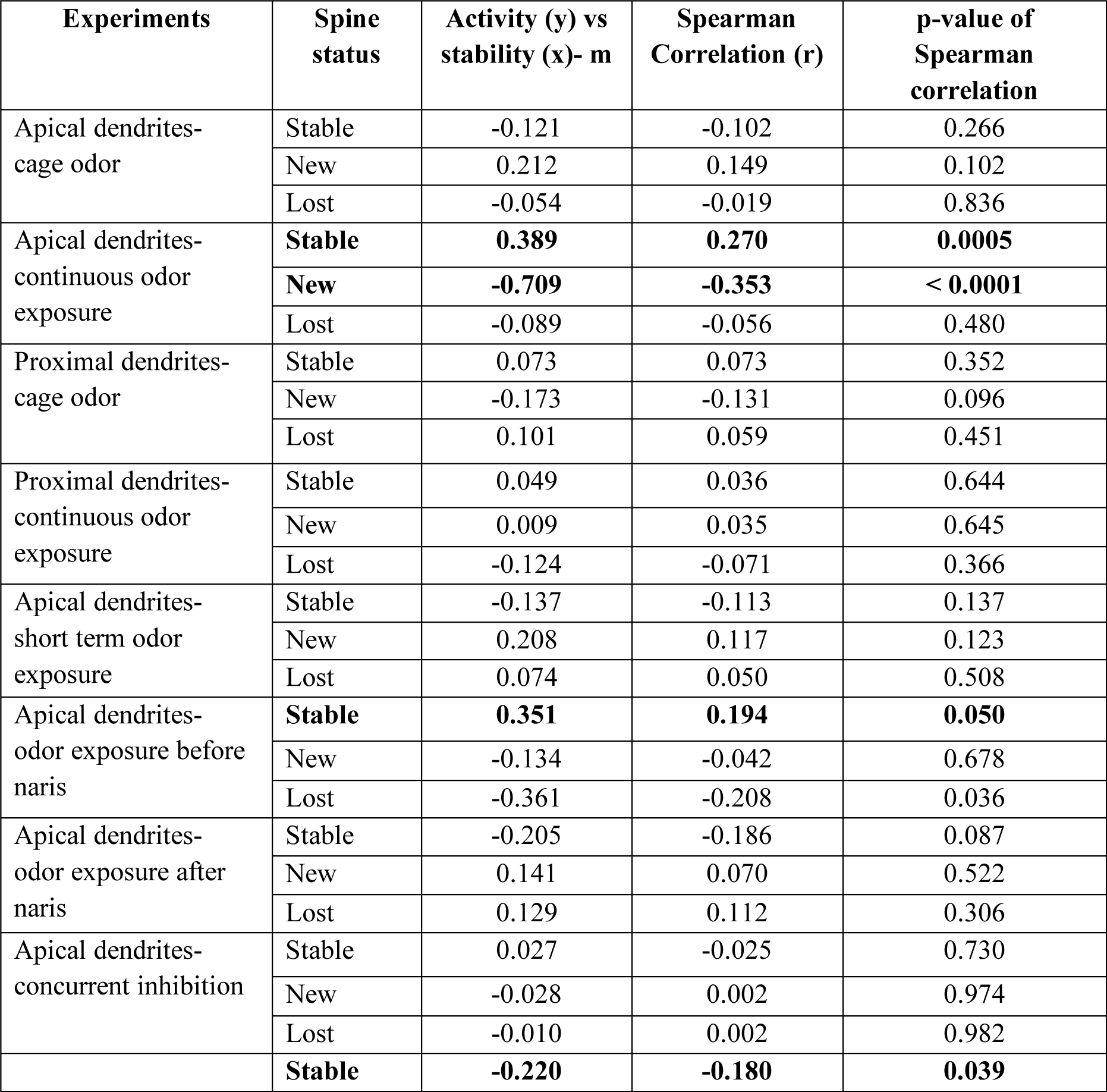
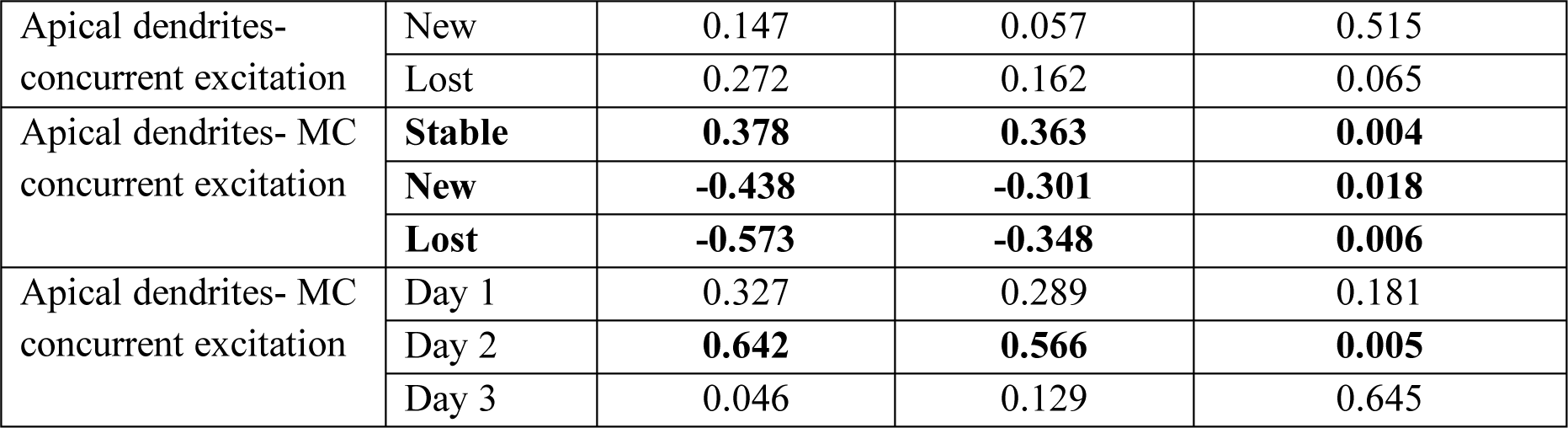
Statistical table for correlations.

**Table 4:** Trends in data with different experiments.

| **Experiments** | **Activity** | **Stability** | **New spines** | **Lost spines** |
| --- | --- | --- | --- | --- |
| **Continuous odor exposure: apical spines** | +++ | + | - | + |
| **Continuous odor exposure: proximal spines** | +++ | +++ | --- | - (ns) |
| **Short term passive odor exposure: apical spines** | +++ | + | -- | - (ns) |
| **Naris occlusion** | --- | - (ns) | -- | ++ |
| **PSD-95: continuous odor exposure** | N/A | + | - (ns) | - (ns) |
| **PSD-95: naris occlusion** | N/A | -- | -- | + |
| **Inhibitory DREADD in GC** | --- | + | --- | + (ns) |
| **Excitatory DREADD in GC** | +++ | + (ns) | - | + (ns) |
| **Excitatory DREADD in MC** | ++ | + (ns) | + | - (ns) |

Increase: +++, $p < 0.0001$ ; ++, $p < 0.01$ , +, $p < 0.05$
Decrease: ---, $p < 0.0001$ ; --, $p < 0.01$ , -, $p < 0.05$

**Table 5:** Values of observations (Mean ± SD)

| Experiments |  | Activity (AUC) | Ampli. | Stable spines | New spines | Lost spines |
| --- | --- | --- | --- | --- | --- | --- |
| Continuous odor exposure: apical spines | Cage odor | 0.256 ± 0.180 | 2.586 ± 1.449 | 0.632 ± 0.164 | 0.176 ± 0.131 | 0.166 ± 0.124 |
| | Odor exposure | 0.390 ± 0.234 | 2.981 ± 1.669 | 0.692 ± 0.157 | 0.148 ± 0.121 | 0.181 ± 0.152 |
| Continuous odor exposure: proximal spines | Cage odor | 0.265 ± 0.116 | 1.841 ± 0.8787 | 0.662 ± 0.135 | 0.191 ± 0.116 | 0.144 ± 0.087 |
| | Odor exposure | 0.431 ± 0.165 | 1.703 ± 0.6416 | 0.767 ± 0.118 | 0.107 ± 0.083 | 0.125 ± 0.081 |
| Short term passive odor exposure: apical spines | Cage odor | 0.323 ± 0.192 | 1.479 ± 1.106 | 0.714 ± 0.153 | 0.155 ± 0.108 | 0.130 ± 0.097 |
| | Passive Odor exposure | 0.599 ± 0.179 | 1.326 ± 0.6534 | 0.763 ± 0.134 | 0.110 ± 0.087 | 0.126 ± 0.109 |
| Naris occlusion | Cage odor | 0.298 ± 0.108 | 1.332 ± 0.5635 | 0.594 ± 0.157 | 0.238 ± 0.132 | 0.167 ± 0.105 |
| | Odor exposure | 0.332 ± 0.147 | 1.53 ± 0.613 | 0.663 ± 0.140 | 0.155 ± 0.093 | 0.181 ± 0.128 |
| | Naris occlusion | 0.126 ± 0.108 | 1.121 ± 0.4172 | 0.558 ± 0.167 | 0.177 ± 0.109 | 0.264 ± 0.176 |
| All spines Td-tomato + PSD-95: | Cage odor | N/A | N/A | 0.722 ± 0.127 | 0.144 ± 0.093 | 0.117 ± 0.073 |
| | Odor exposure | N/A | N/A | 0.796 ± 0.094 | 0.092 ± 0.074 | 0.122 ± 0.084 |
|  | Naris occlusion | N/A | N/A | 0.716 ± 0.125 | 0.122 ± 0.087 | 0.161 ± 0.112 |
| PSD-95+ spines only | Cage odor | N/A | N/A | 0.588 ± 0.229 | 0.238 ± 0.169 | 0.213 ± 0.160 |
|  | Odor exposure | N/A | N/A | 0.635 ± 0.229 | 0.221 ± 0.162 | 0.192 ± 0.175 |
|  | Naris occlusion | N/A | N/A | 0.574 ± 0.234 | 0.141 ± 0.151 | 0.284 ± 0.202 |
| Inhibitory DREADD in GC | Cage odor | 0.298 ± 0.135 | 1.715 ± 0.680 | 0.656 ± 0.128 | 0.201 ± 0.114 | 0.142 ± 0.081 |
|  | After CNO | 0.191 ± 0.096 | 1.376 ± 0.645 | 0.688 ± 0.113 | 0.150 ± 0.094 | 0.162 ± 0.091 |
| Excitatory DREADD in GC | Cage odor | 0.352 ± 0.162 | 1.254 ± 0.527 | 0.588 ± 0.181 | 0.247 ± 0.137 | 0.201 ± 0.121 |
|  | After CNO | 0.492 ± 0.164 | 1.434 ± 0.607 | 0.584 ± 0.149 | 0.206 ± 0.109 | 0.212 ± 0.117 |
| Excitatory DREADD in MC | Cage odor | 0.254 ± 0.143 | 1.325 ± 0.653 | 0.592 ± 0.176 | 0.249 ± 0.157 | 0.147 ± 0.112 |
|  | After CNO | 0.391 ± 0.222 | 1.214 ± 0.546 | 0.653 ± 0.206 | 0.160 ± 0.115 | 0.177 ± 0.119 |

#### Power test

Statistical power analysis was performed using G*Power version 3 ^52^ and reported in Table 2.

### A two-stage model of structural plasticity in the OB

The firing rates of MCs and GCs are described by the ordinary differential equations ^10^,

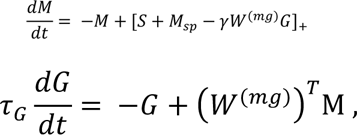

where the firing rates *M* and *G* are vectors of size *NMC* and *NGC*, respectively, []+ denotes the piecewise linear rectifier, and γ gives the inhibitory strength. The MCs receive sensory inputs *S* and have spontaneous activity *M*_*sp*_. For steady input and fixed connectivity *W^(mg)^* the neuronal activities always converge to a steady state. In the following, neuronal activities always refer to this steady state.

The reciprocal connectivity between the MCs and GCs is given by the connectivity matrix *W^(mg)^* and its transpose (*W*^(*mg*)^V^*T*^. In the OB the connections are via dendrodendritic synapses located on the secondary dendrites of the MCs, which reach across large portions of the bulb, but only sparsely so. We mimic the resulting geometric constraint on the connectivity by allowing each GC to connect only to a randomly chosen set of MCs. The focus of this model is the structural plasticity of the synapses, which leads to an activity-dependent evolution of *W^(mg)^*.

Since not all synapses included in the experimental stability statistics contain PSD-95 (Fig.6), we consider two types of synapses: nonfunctional and functional ones. We assume that the formation of a fully functional synapse, which may be identified with the spines expressing PSD-95 in addition to Td-Tomato in Supp. Fig.6, occurs through a consolidation process of a non-functional spine that only expresses Td-Tomato. This consolidation may be related to the formation of the postsynaptic density.

We take the formation of a nonfunctional synapse between an MC and a GC to have a probability that depends on the activity of that MC, while the removal rate is assumed to be activity-independent,

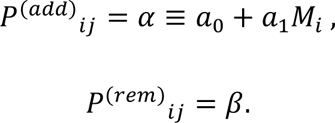

The activity-dependence of the formation is motivated by the dependence of the formation of filopodia on glutamate and BDNF released by stimulated MCs as reported ^18, 26^ . The presence of nonfunctional synapses is stored in a matrix *ŵ*_*ij*_^(*mg*)^.

The consolidation of a nonfunctional synapse and the deconsolidation of a functional synapse are assumed to depend on the firing rates *M*_*i*_ and *G*_*j*_ of the MC *i* and of the GC *j*, respectively, that are connected by that synapse and on the size *P*_j_^(*psd*)^ of a pool of resources on the GC needed for the consolidation. We express the consolidation and deconsolidation rates in terms of a single rate function (cf. Fig.7b),

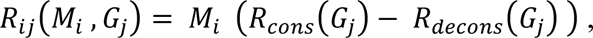

where *Rcons* and *Rdecons* are two sigmoidal rate functions,

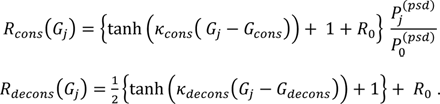

Here *k*_*cons*_ and *k*_*decons*_ are the slopes at the inflection points *G*_*cons*_ and *G*_*decons*_ of the sigmoidal functions respectively. Note that even for *G*_*j*_ = 0 some consolidation and deconsolidation is going on depending on *R*_0_ and *P*_*j*_^(*psd*)^.

The consolidation rate increases with GC activity, which in turn increases with an increase in the number of consolidated synapses. To avoid a run-away formation of consolidated synapses through this positive feedback, we assume that the consolidation depletes the resource pool *P^(psd)^*. Performing a time step of size Δt, the available pool *P*_*j*_^(*psd*)^ on GC *j* evolves therefore according to

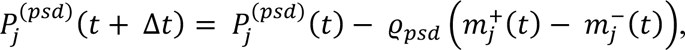

where *mj*^+^ (*t*) and *mj*^+^ (*t*) are the number of functional spines that are being consolidated and deconsolidated, respectively, and *ϱpsd* is the resource amount required for each consolidation. Effectively, the depletion of the resource pool shifts the threshold for the consolidation process upward, reminiscent of the sliding threshold in the BCM model for synaptic weight plasticity^53^. The role of the resource pool is illustrated in Fig.7b, where synapses (red) are filled (blue) when they are consolidated. MC activity that drives the GC above threshold (red arrow) consolidates the synapse and reduces the resource pool, while MC activity that does not drive the GC sufficiently to render R positive (orange arrow), deconsolidates the synapse and refills the pool. Inputs that drive the GC very weakly (blue arrow) do not change the synapse.

Specifically, in a given time step of size *Δt*, an unconsolidated synapse becomes consolidated with probability

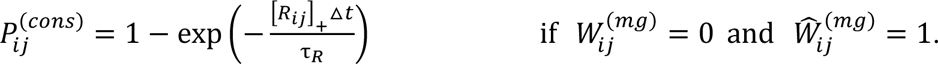

Conversely, a consolidated synapse is deconsolidated with probability

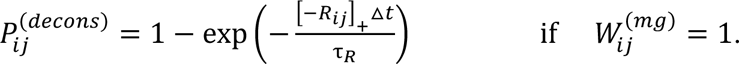

We allow heterogeneity in the plasticity parameters Gcons and Gdecons across the GCs and set for each 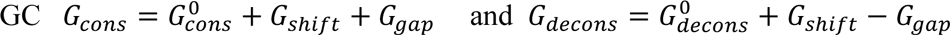, where *G*_*shift*_ and *G*_*gap*_ are uniformly distributed within the ranges *G*_*shift*_ and *G*_*gap*_, respectively.

As initial condition we start each simulation with each GC having 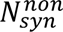 nonfunctional (unconsolidated) synapses and 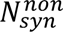 functional synapses within its geometrically allowed connection set of *N*_*conn*_ synapses.

The stimuli are based on the glomerular activity patterns of Leon/Johnson ^27^. To speed up the computations we have down-sampled them to 236 input channels corresponding to 236 MCs. Since these activity patterns are given in terms of a z-score, their means are not included in the data. Experimentally, it is found that typically on the order of 30% of the MCs are active for a given stimulus ^21^. We therefore rescale the stimuli with a threshold *S*_*thr*_ chosen such that about 30% of the MCs are excited,

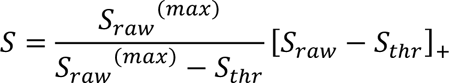

To generate stimuli that are difficult to discriminate, we use mixtures through linear composition of the individual components. The resulting stimulus patterns that were used in the computations are shown in Fig.7E.

The model is implemented in Matlab. The parameters are given in the table.

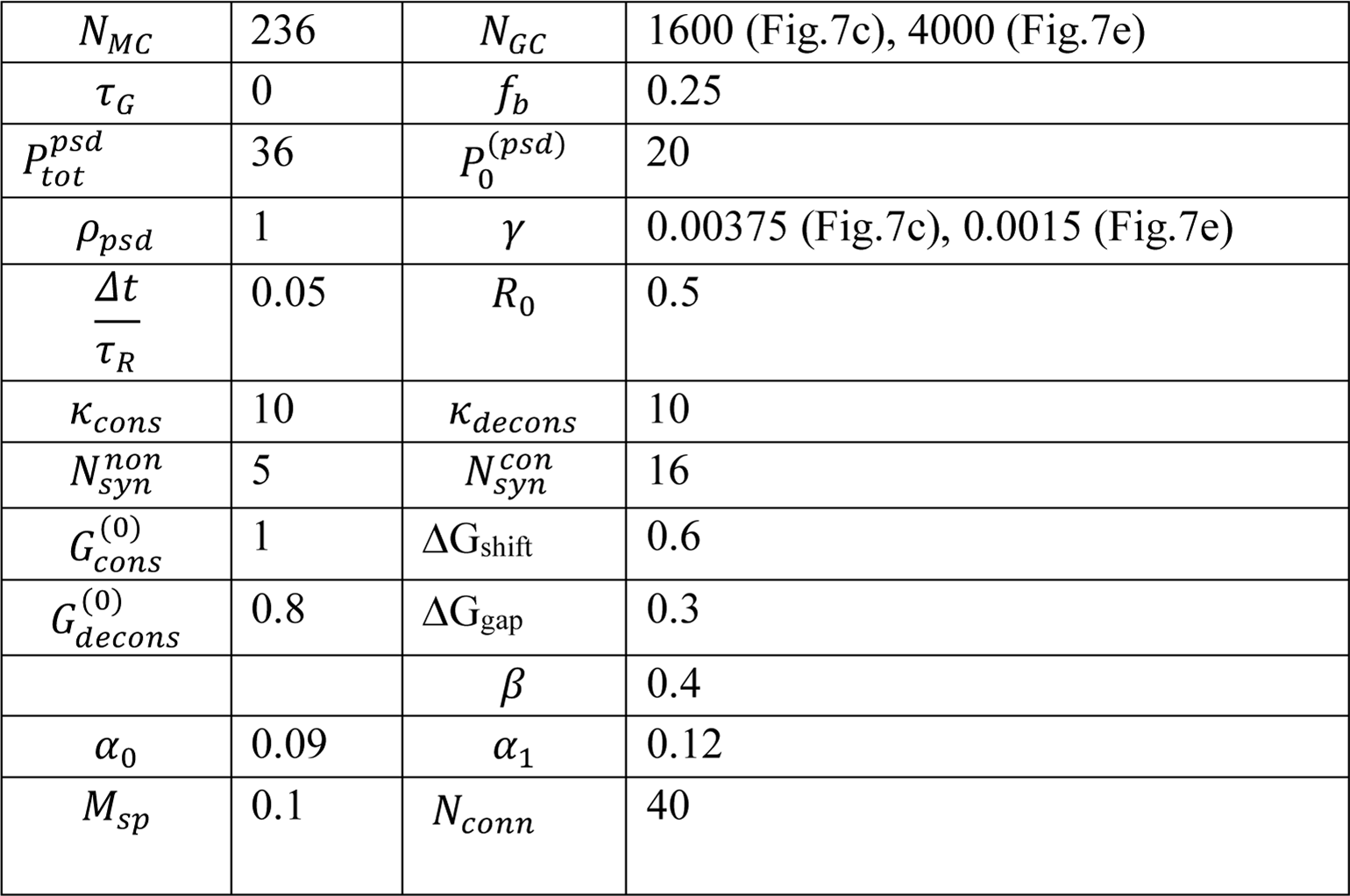

|  |  |  |  |
| --- | --- | --- | --- |
| $N_{MC}$ | 236 | $N_{GC}$ | 1600 (Fig.7c), 4000 (Fig.7e) |
| $\tau_G$ | 0 | $f_b$ | 0.25 |
| $P_{tot}^{psd}$ | 36 | $P_0^{(psd)}$ | 20 |
| $\rho_{psd}$ | 1 | $\gamma$ | 0.00375 (Fig.7c), 0.0015 (Fig.7e) |
| $\frac{\Delta t}{\tau_R}$ | 0.05 | $R_0$ | 0.5 |
| $\kappa_{cons}$ | 10 | $\kappa_{decons}$ | 10 |
| $N_{syn}^{non}$ | 5 | $N_{syn}^{con}$ | 16 |
| $G_{cons}^{(0)}$ | 1 | $\Delta G_{shift}$ | 0.6 |
| $G_{decons}^{(0)}$ | 0.8 | $\Delta G_{gap}$ | 0.3 |
| | | $\beta$ | 0.4 |
| $\alpha_0$ | 0.09 | $\alpha_1$ | 0.12 |
| $M_{sp}$ | 0.1 | $N_{conn}$ | 40 |

We assessed the discriminability of pairs of activity patterns *M*^(1,2)^ using the Fisher discriminant, which compares the difference in the means of the patterns with their trial-to-trial variability. Since the firing rate model does not include such variability, we assumed that the firing rates arise from Poisson spike trains. The variance of the spike counts is then given by their means. Assuming a linear read-out of the activity patterns with a weight vector w that maximizes the Fisher discriminant, the Fisher discriminant for the read-outs of the two activity patterns is then given by

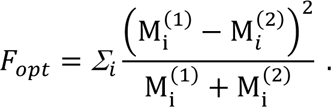

## Extended Data information

**Extended Data Figure 1:**
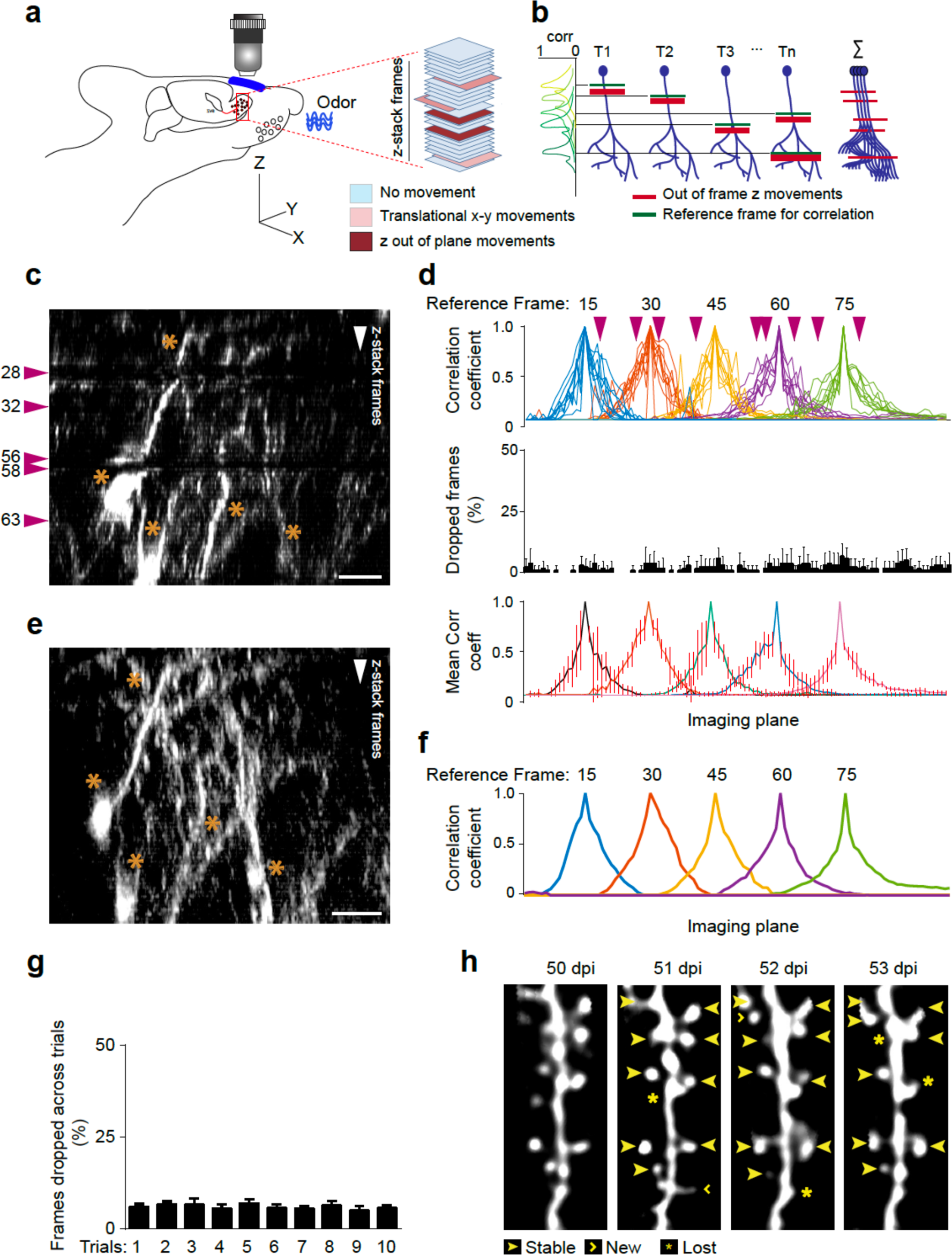
Protocol for structural in vivo 2-photon imaging in awake mice. a. Schematic representation of *in vivo* imaging of 3-D structural stacks in awake mice, highlighting the principle types of movements encountered during imaging. The magnification in red represents a representative z-stack where two types of movements are highlighted: x-y lateral frame movements (light red color) and out-of-plane z-movements (in dark red). b. Representative example of 2-D cross correlation technique to eliminate the out-of-plane z-movements from a neuronal structural image. T1…Tn indicate trial numbers of fast-imaging on the same field of view using the resonant galvanometer scanner. Red horizontal bars indicate ‘actual’ frames where movement has taken place during that trial while green represents a ‘reference’ frame against which a 2-D cross correlation is performed across all the frames in that trial. The correlation plot on the left demonstrates the correlation plot of the frames, and ‘breaks’ seen in the distribution which determines the actual frame of movement. Once determined, these frames are replaced with ‘NaN’ values and averaged across all the trials using *nanmean* function in MATLAB. c. Example from an imaging experiment in the z-plane which shows the loss of information due to movements. The closed arrows indicate the points of movement in the raw image during one of the trials. Scale bar = 15um. Asterisks (*) indicates the structures in the image used for later comparison. d. Correlation coefficient plot showing the values of Pearson’s r with respect to the reference frames 15, 30, 45, 60 and 75 (15 frame interval) selected from the z-stacks and the whole z-stack across the multiple trials of fast imaging. The correlation plots are fitted with an ideal distribution with maximum correlation at the reference frames, and points where they diverge from the ideal plot are noted. These points correspond to out-of-z frame movements. Those points are indexed and replaced by ‘NaN’ values. The second panel demonstrates the percentage of frames dropped across all the trials. The third panel shows the mean correlation coefficient plot across the trials. The error bars indicate standard deviation. e. Example of reconstruction of the same z-stack as in C, with the dropped frames replaced by ‘NaN’ and averaged across the trials for the particular region of imaging. Note that the structure reconstructed have no loss of structural information owing to elimination of some z-frames in some trials. Asterisks (*) demonstrate portions that are able to recount for lost structures shown in C. f. Correlation coefficient plot of the reconstructed image with respect to reference frames at an interval of 15 frames. Note that the movement artefacts are removed compared to the correlation coefficient plot as in D, first panel. g. Quantification of the percentage of frames dropped across 7 animals across 10 trials of the awake structural imaging reconstructions. h. Example 2-photon projected image of a dendritic segment tracked across 4 days (50dpi – 53dpi) of awake structural imaging. The new, stable and lost spines along the dendrites are indicated according to the convention as shown (closed arrow: stable spine, open arrow: dynamic spines and star: lost spines).

**Extended Data Figure 2:**
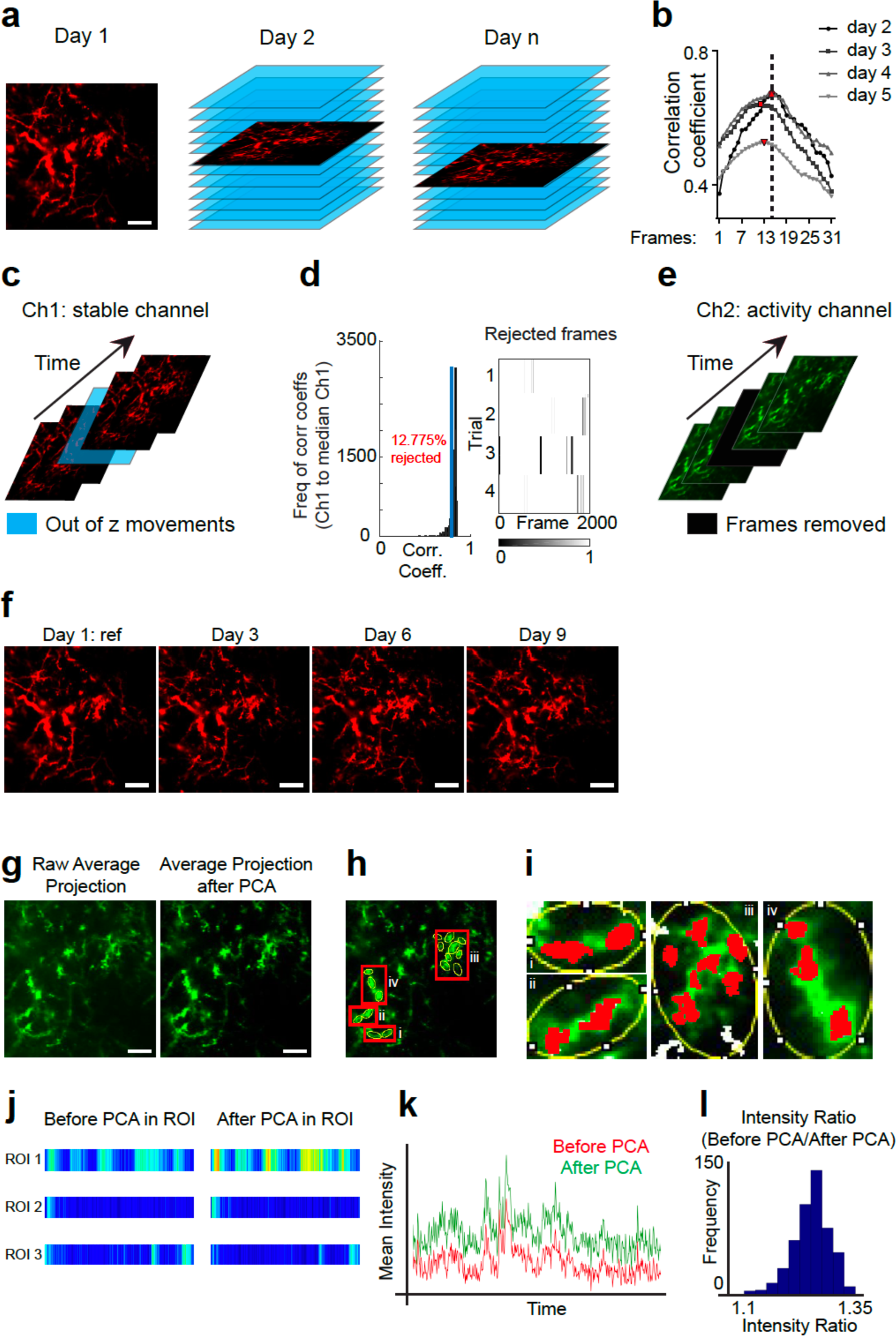
Protocol for neuronal activity using *in vivo* 2-photon imaging in awake mice. a. Schematic representation demonstrating the identification of the same plane of imaging across multiple days to record neuronal activity from the same region of imaging. The average projection of the stable channel (Ch1, Td tomato) was used in a time series recording from a plane of imaging selected on day 1 and use as the reference for subsequent days of imaging. On the following days, the estimated z-plane of imaging was manually found and the day 1 z-plane reference was used to match the correct z-plane frame using a 2-D cross correlation of the reference frame to the z-stack taken on subsequent days. b. Values of cross-correlation of the reference frame to the example z-stack taken on subsequent days. The dotted vertical line indicates the human estimation of the day 1 reference frame while red spots indicate the frames with the highest 2-D correlation coefficient in the corresponding days of imaging. These indices are used to select the imaging plane for recording activity across multiple days. c. Schematic example of out-of-z-frame movement artefact during imaging of neuronal activity. Using the tdTomato signal (in red, Ch1: stable channel), the out of frame movement was detected in time, as indicated by the blue frame. d. Histogram plot of the 2-D correlation coefficients between the average projection of Ch1 across the entire time series with each Ch1 frame in the trial. The blue vertical line indicates the threshold of selection for rejected frames. The rejected frame plot indicates the frames in the trial which are below the selected threshold and rejected (black vertical lines). These frames of the trial are indexed and excluded from analysis. e. Schematic example showing the removal of corresponding frames on GCaMP6f recording (in green, Ch2: activity channel) based on the indexed frames from stable channel Td tomato (in d). The removed frames are replaced with ‘NaN’ values to preserve the indices of the removed frames and these frames are excluded from analysis. f. Day to day registration protocol: The average projection of the stable channel (Ch1, Td tomato) across each day was taken and used for registration to correct for x-y lateral movements for each dat. The Day 1 average projection was taken as the reference. The registration values were calculated in ImageJ (*Multistackreg*), and then propagated to all frames in the activity channel (Ch 2, GCaMP6f). g. Post processing of motion corrected activity time series: Removal of noise by using principal component analysis (PCA)-assisted reconstruction of the raw image pixels. Raw image pixels were vectorized and transposed into eigen vectors to isolate orthogonal vectors adding to maximum variance of signals across the dataset, and reconstructed using the first 25 principal components (PCs). Average projection after PCA indicates significant reduction of noise from the data as compared to the raw average projection. h. Selection of regions of interest (ROIs) after the PCA-assisted reconstruction. i. Example ROIs (expanded from I) show the redefined boundary of the ROIs after reconstructing each ROIs using the 1^st^ PC. Note that the new boundary only encompasses pixels smaller than the original elliptical ROIs. j. Example of noise reduction and signal-to-noise enhancement before and after PCA in three example ROIs. k. Plot showing the improvement of signal-to-noise in pixel values obtained from a ROI across time after PCA reconstruction (green) compared to before PCA reconstruction (red). l. Frequency distribution of signal enhancement after PCA reconstruction. The median value is 1.25 indicating a 25% improvement in the signal to noise ratio.

**Extended Data Figure 3:**
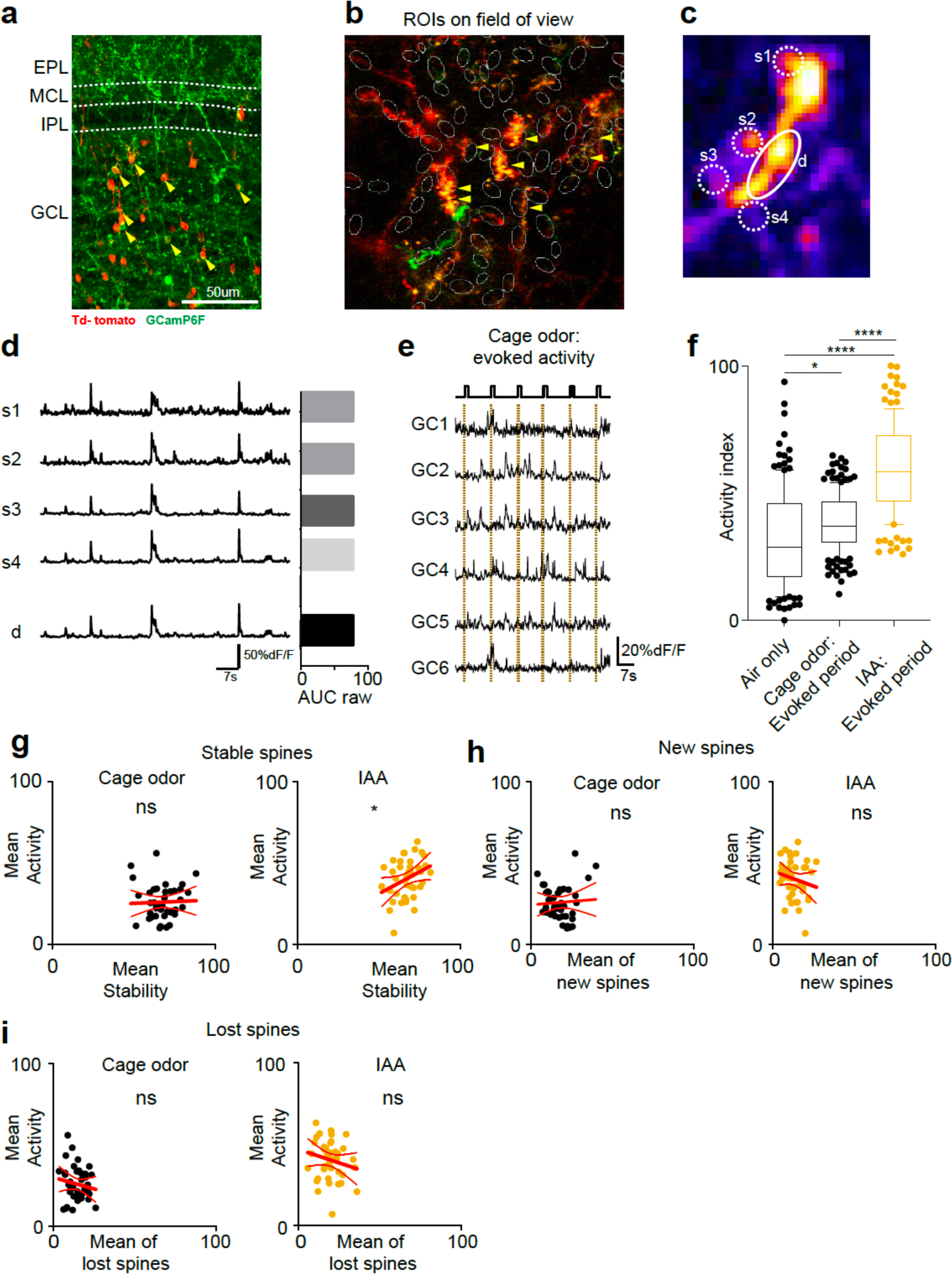
GC labelling techniques, ROI selection, day-to-day spine dynamics and neuronal activity. a. Confocal image of different layers of the olfactory bulb (OB) showing cre-recombinase virus expressing tdTomato (red) and their co-localization with floxed GCaMP6f (green) in GC within the GCL (arrows). Scale bar = 50um. b. Merged average projection of a field of view under 2-photon microscope showing GC dendrites co-expressing tdTomato and GCaMP6f. The ROIs are selected on the dendrites expressing both the markers. c. Merged average projection of a GCaMP6f-expressing GC dendrite and spines under 2-photon microscope. s1-4 are the ROIs on the spines and ‘d’ is an ROI on the dendritic segment. d. Example Ca^2+^ traces from the ROIs selected in c, along with the raw magnitude of their activity in terms of AUC. Scale: 50%dF/F, 7s. (Right) Raw activity (Area under the curve) quantification in example ROIs as indicated. e. Example traces of GCs in response to evoked exposure to cage odor (2s). The rectangle at the top indicates the odor delivery times during the activity recording. f. Boxplots showing the min-max normalized neuronal activity in air only and odor evoked condition of cage odor and IAA in ROI-dendrite pairs (black: air only and cage odor background, gold: IAA exposure). g. Scatter plot of mean neuronal activity (activity index) versus mean percentage of stable spines in cage odor background and continuous IAA with linear regression fit. h. Scatter plot of mean neuronal activity (activity index) versus mean percentage of new spines in cage odor background and continuous IAA with linear regression fit. i. Scatter plot of mean neuronal activity (activity index) versus mean percentage of lost spines in cage odor background and continuous IAA with linear regression fit.

**Extended Data Figure 4:**
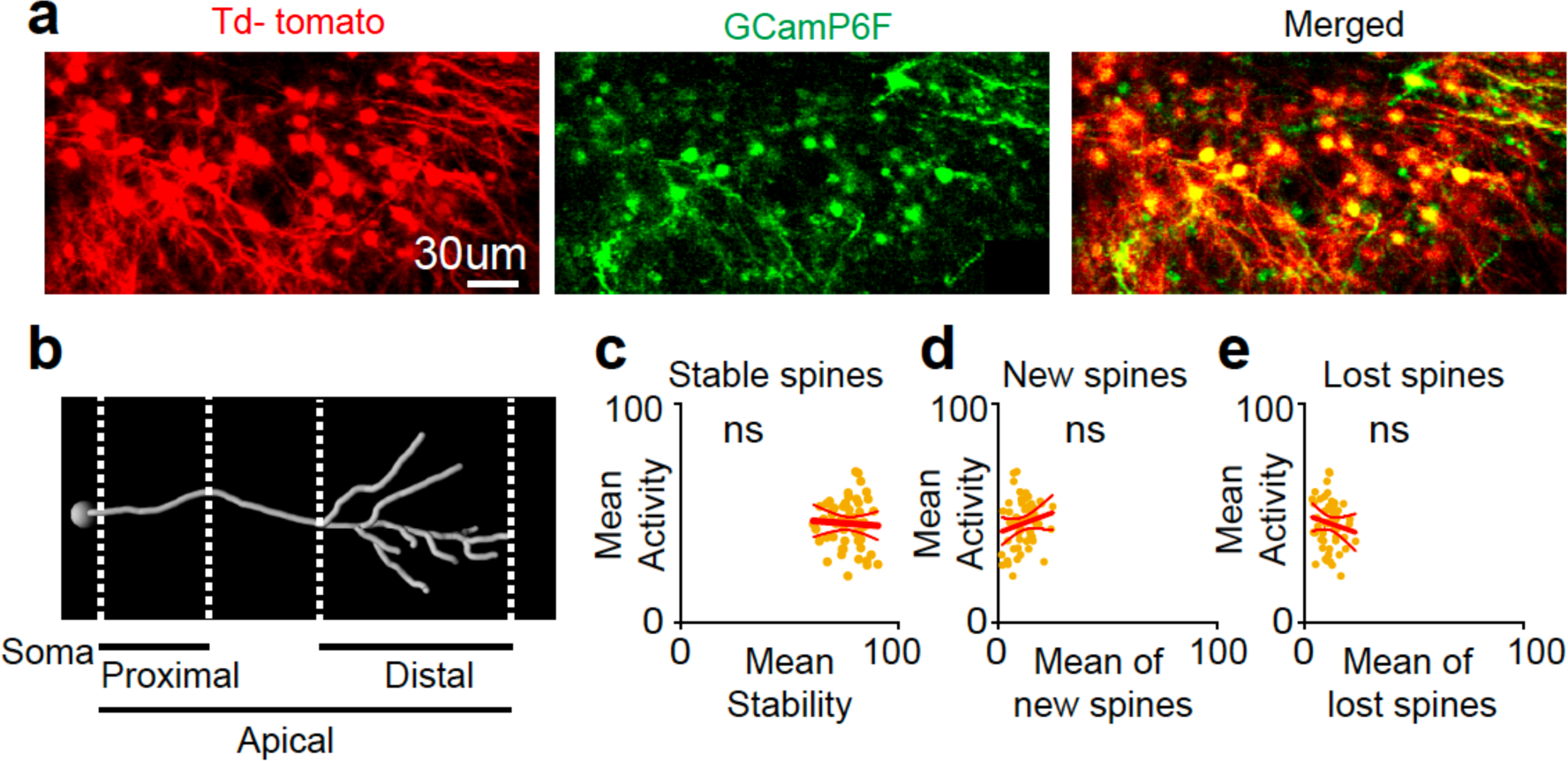
Proximal GC dendrite recordings, day-to-day spine dynamics and neuronal activity. a. Average projection of a field of view for imaging the proximal dendritic regions of GCs. b. A tracing reconstruction of a GC demonstrating the different dendritic segments. The proximal region of the dendrite used in this experiment, is indicated. c. Scatter plot of mean neuronal activity (activity index) versus mean percentage of stable spines in continuous IAA with linear regression fit. d. Scatter plot of mean neuronal activity (activity index) versus mean percentage of new spines in continuous IAA with linear regression fit. e. Scatter plot of mean neuronal activity (activity index) versus mean percentage of lost spines in continuous IAA with linear regression fit.

**Extended Data Figure 5:**
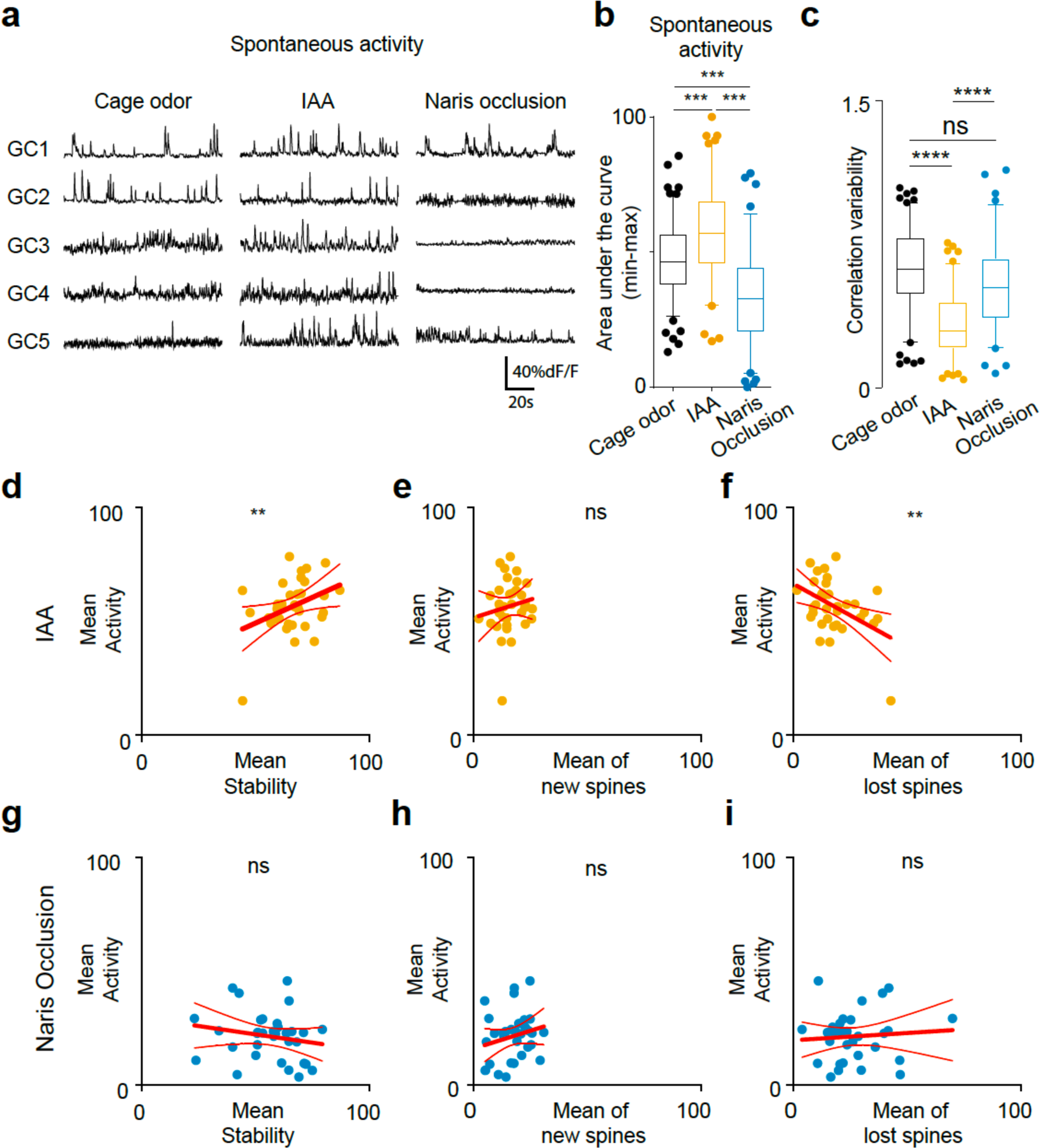
Apical GC dendrite recordings in naris occlusion, day-to-day spine dynamics and neuronal activity. a. Example Ca^2+^ traces of GC spontaneous activity in cage odor, continuous IAA and IAA with naris occlusion. b. Boxplots showing the distribution of the min-max normalized spontaneous activity of the ROI-dendrite pairs for three experimental conditions: cage odor, continuous IAA and IAA with naris occlusion. c. Boxplot of the correlation variability of the individual dendrite-ROIs from their mean for cage odor background (black), continuous IAA (gold) and naris occlusion (blue) shows a significant decrease for IAA condition while significant increase for naris occlusion compared to IAA. d. Scatter plot of mean neuronal activity (activity index) versus mean percentage of stable spines in continuous IAA with linear regression fit. e. Scatter plot of mean neuronal activity (activity index) versus mean percentage of new spines in continuous IAA with linear regression fit. f. Scatter plot of mean neuronal activity (activity index) versus mean percentage of lost spines in continuous IAA with linear regression fit. g. Scatter plot of mean neuronal activity (activity index) versus mean percentage of stable spines in naris occlusion with linear regression fit. h. Scatter plot of mean neuronal activity (activity index) versus mean percentage of new spines in naris occlusion with linear regression fit. i. Scatter plot of mean neuronal activity (activity index) versus mean percentage of lost spines in naris occlusion with linear regression fit.

**Extended Data Figure 6:**
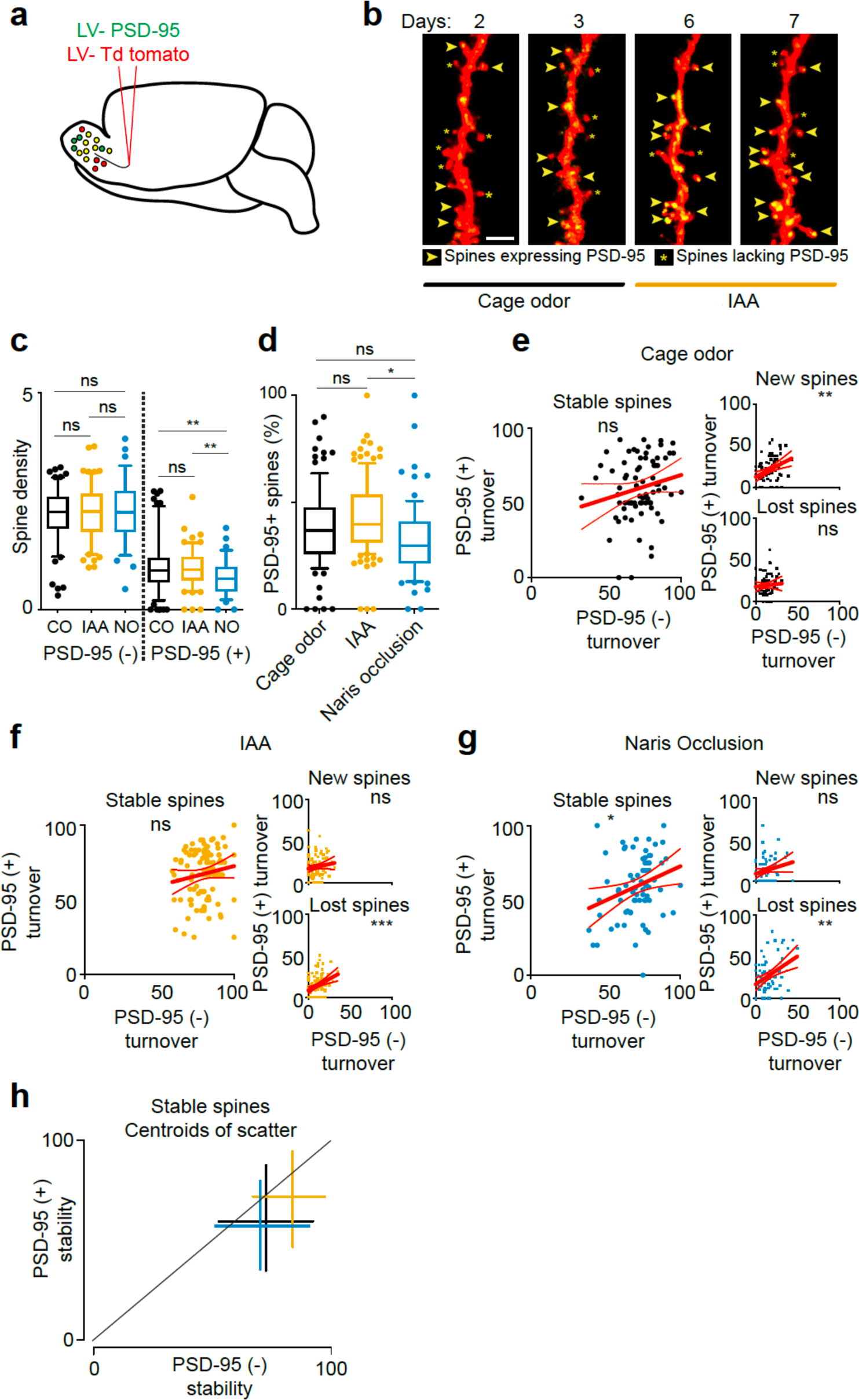
Naris occlusion; PSD-95 turnover. a. Viral labelling strategy to co-label adult born GCs with lentiviral capsids containing PSD95-GFP and tdTomato reporters. b. 2-photon projected images from day 2 to 3 and 6 to 7, corresponding to cage odor and IAA exposure, respectively. Spines expressing or lacking PSD-95 are indicated in the figure. Scale: 5um. c. Boxplot showing the spine density across the dendrites for all spines (expressing PSD 95 and tdTomato, left) and spines expressing PSD-95 exclusively (right). d. Box plot showing the percentage of spines expressing PSD-95 in GCs. With continuous IAA, the percentage of PSD-95^+^ spines increase (not significant to cage odor), and in naris occlusion, there is a significant decrease of PSD-95^+^ spines. It is to be noted that not all spines express PSD-95. e. Scatter plot of stable, new and lost spine turnover for PSD-95^+^ and PSD-95^-^ spines in cage odor condition with linear regression fit. f. Scatter plot of stable, new and lost spine turnover for PSD-95^+^ and PSD-95^-^ spines in continuous IAA with linear regression fit. g. Scatter plot of stable, new and lost spine turnover for PSD-95^+^ and PSD-95^-^ spines in naris occlusion with linear regression fit. h. Centroids of the scatter plots of the stable spines (f-h) with the standard deviation.

**Extended Data Figure 7:**
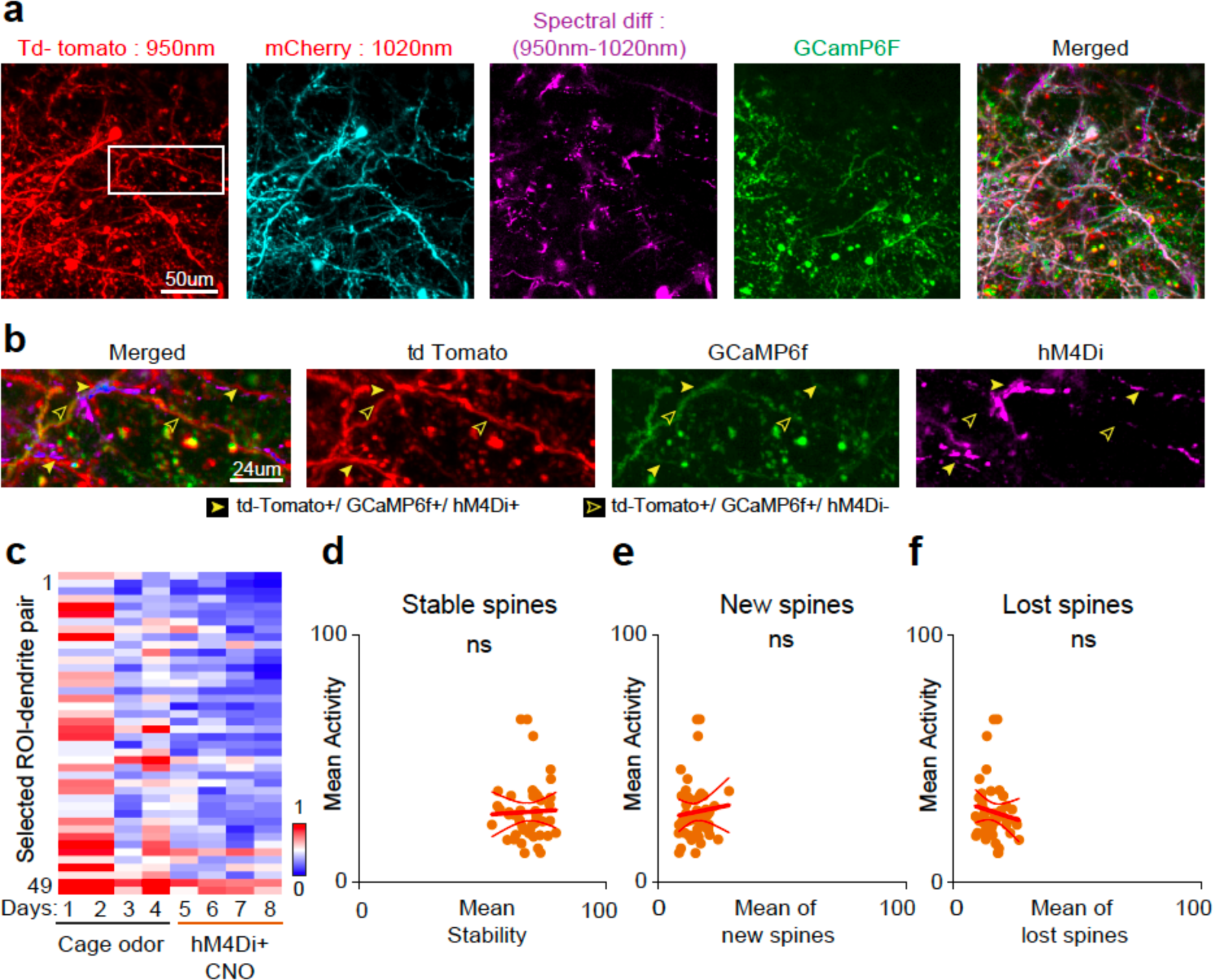
Apical GC dendrite recordings in chemo-genetic inhibition, day to day spine dynamics and neuronal activity. a. Average intensity projection of a field of view at wavelength 950nm (for tdTomato expression), 1020nm (for mCherry expression), spectral difference to isolate mCherry expression (hM4Di; difference of 950nm and 1020nm) and GCaMP6f. The merged photomicrograph represents the expression of tdTomato, GCaMP6f and the spectral difference (hM4Di). b. Representative image demonstrating selection of ROIs based on the criteria of tdTomato^+^, GCaMP6f^+^ and hM4Di^+^expressions. c. Heat map showing the neuronal activity (AUC, min-max normalized across the dataset) averaged per day across different experiments (49 dendrite-ROI pairs, 5 animals). d. Scatter plot of mean neuronal activity (activity index) versus mean percentage of stable spines in CNO-mediated inhibition with linear regression fit. e. Scatter plot of mean neuronal activity (activity index) versus mean percentage of new spines in CNO-mediated inhibition with linear regression fit. f. Scatter plot of mean neuronal activity (activity index) versus mean percentage of lost spines in CNO-mediated inhibition with linear regression fit.

**Extended Data Figure 8:**
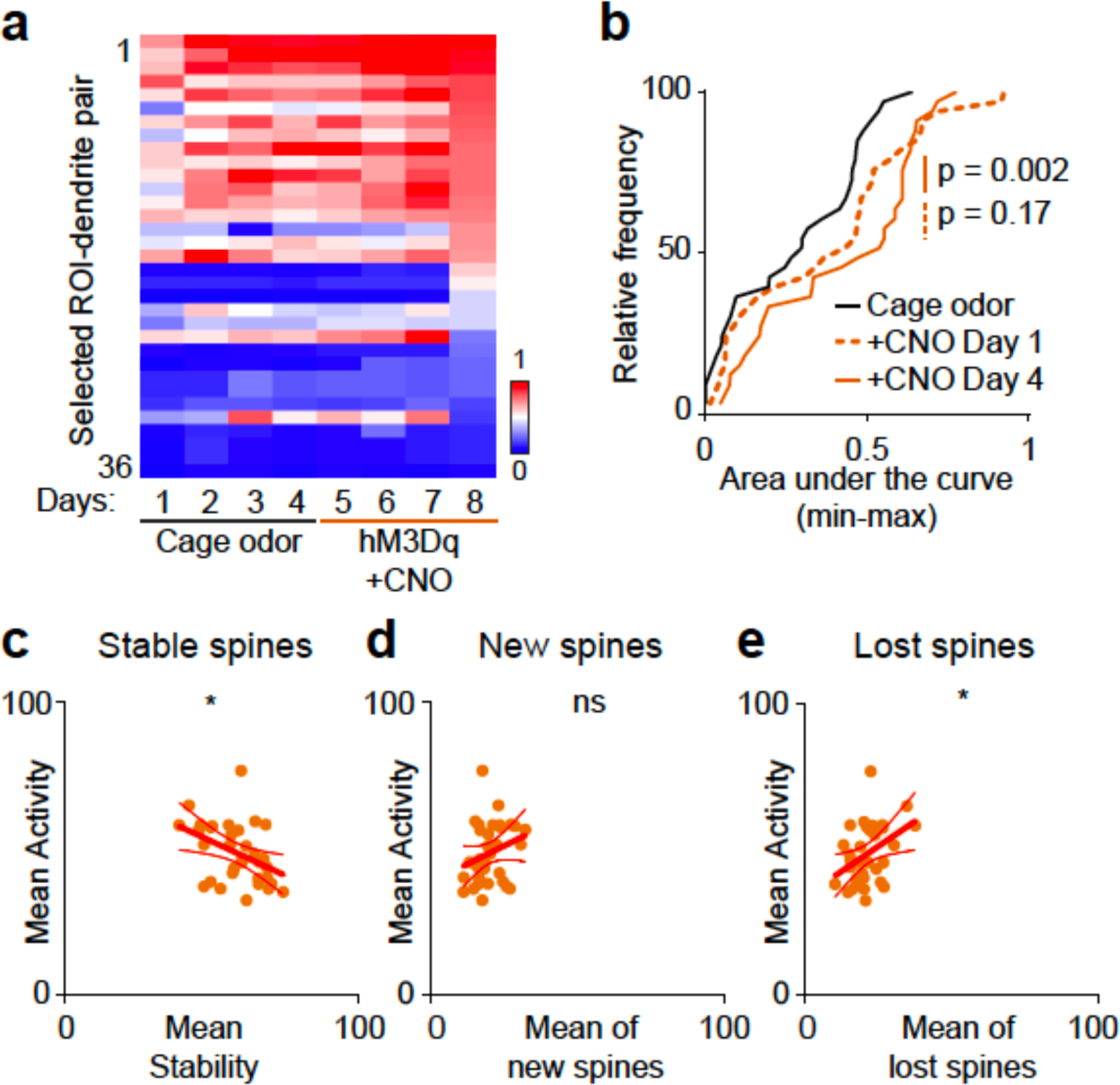
Chemo-genetic excitation, day to day spine dynamics and neuronal activity. a. Heat map showing the neuronal activity (AUC, min-max normalized across the dataset) averaged per day across different experiments (36 dendrite-ROI pairs, 4 animals). b. Relative frequency plot of neuronal activity showing the effect of chronic CNO-mediated GC excitation between cage odor on day 1 of CNO application (p = 0.17) and on day 4 of CNO application. c. Scatter plot of mean neuronal activity (activity index) versus mean percentage of stable spines in CNO-mediated excitation with linear regression fit. d. Scatter plot of mean neuronal activity (activity index) versus mean percentage of new spines in CNO-mediated excitation with linear regression fit. e. Scatter plot of mean neuronal activity (activity index) versus mean percentage of lost spines in CNO-mediated excitation with linear regression fit.

**Extended Data Figure 9:**
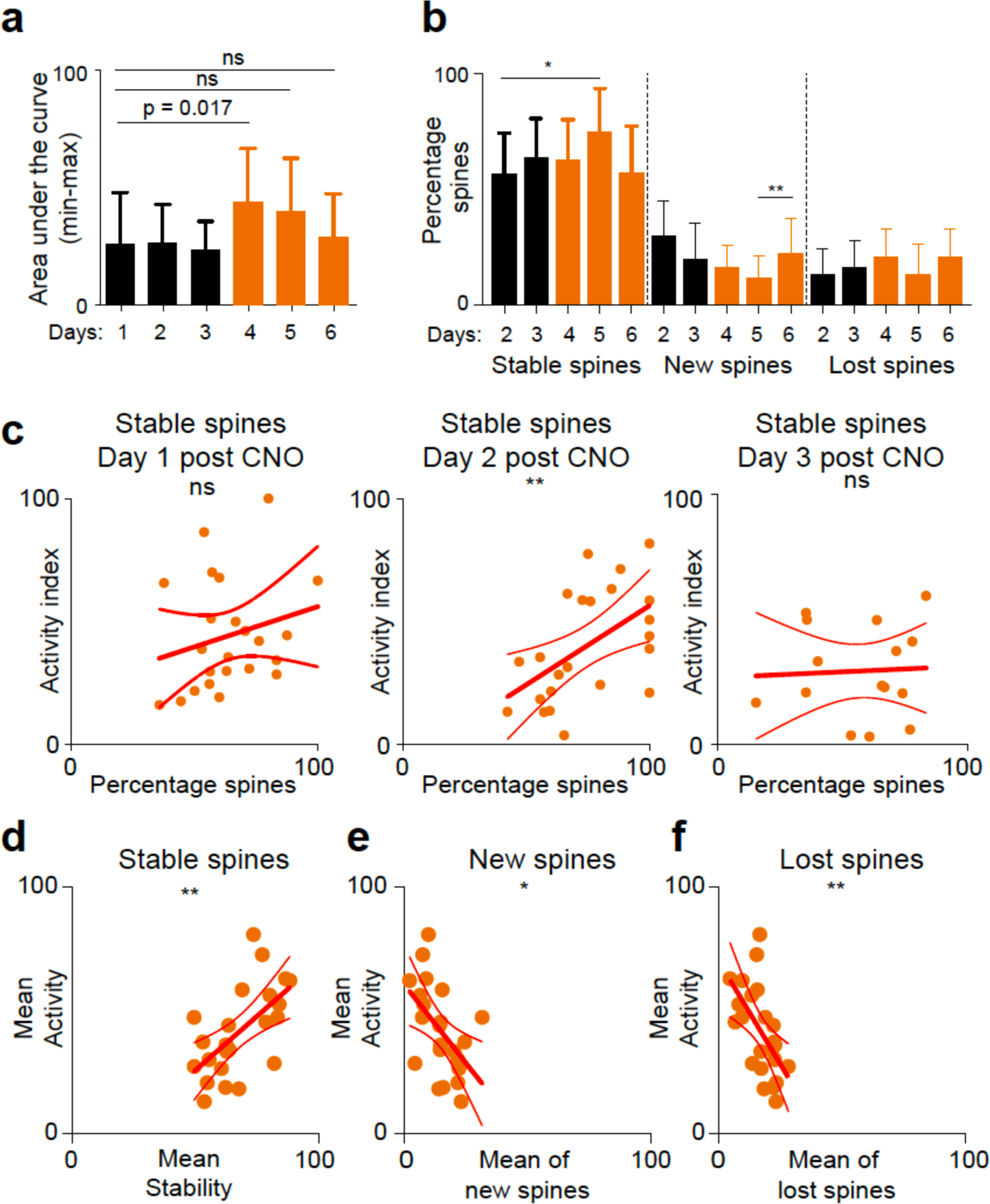
Additional data on MC DREADD activation. a. Bar-plots showing mean neuronal activity of GCs across days in selected ROI-dendrite pairs across different experiments (black: cage odor background, red: CNO-mediated MC activation). b. Percentage turnover of stable, new and lost spines in the selected ROI-dendrite pairs (black: cage odor background; red: CNO-mediated MC activation) at one day interval. c. Scatter plot of neuronal activity (activity index) versus percentage of stable spines in chronic CNO-mediated MC excitation with a linear regression fit across days 1-3 after CNO application. d. Scatter plot of mean neuronal activity (activity index) versus mean percentage of stable spines in CNO-mediated MC excitation with linear regression fit. e. Scatter plot of mean neuronal activity (activity index) versus mean percentage of new spines in CNO-mediated MC excitation with linear regression fit. f. Scatter plot of mean neuronal activity (activity index) versus mean percentage of lost spines in CNO-mediated MC excitation with linear regression fit.

**Extended Data Figure 10:**
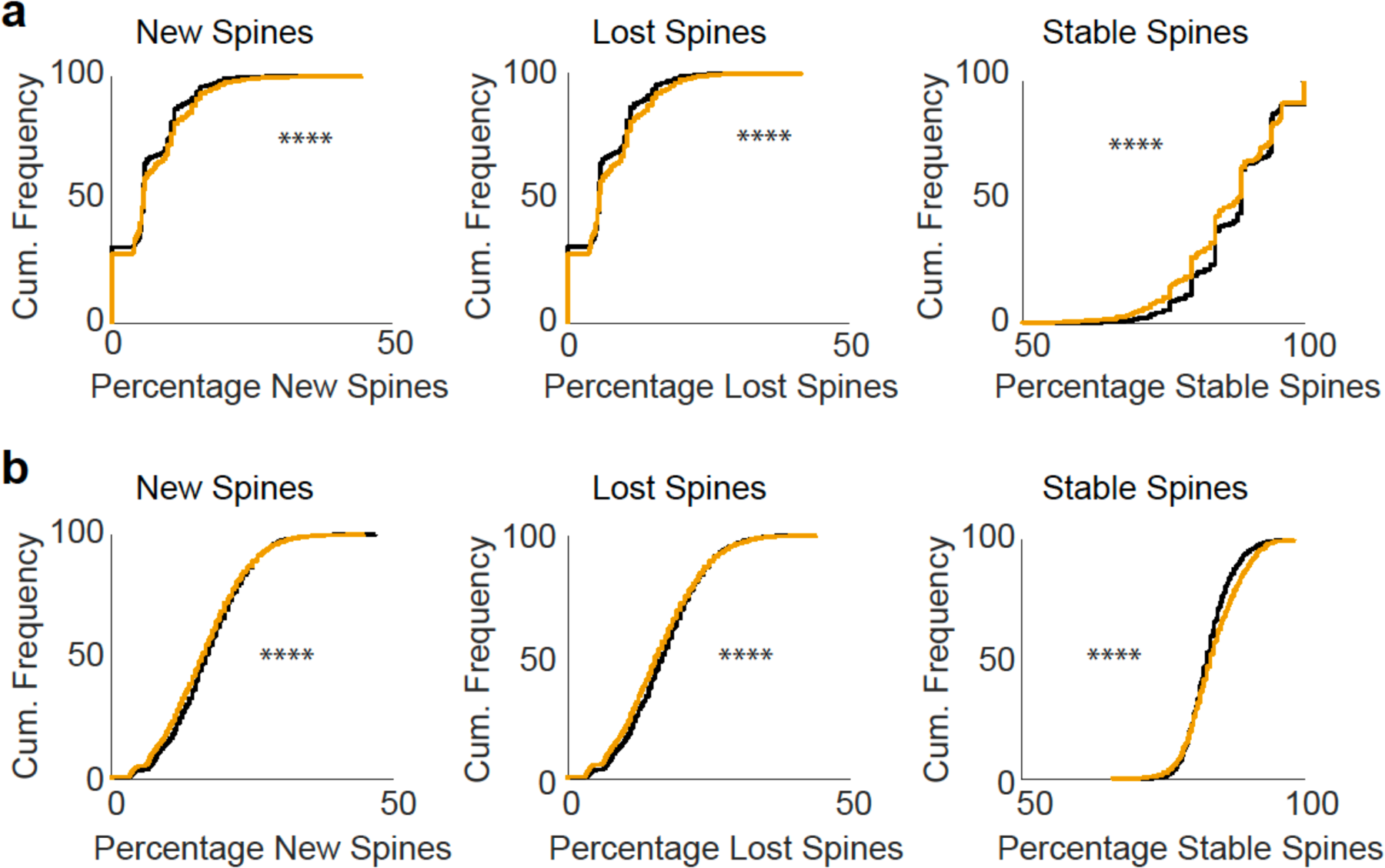
Dependence of spine formation on MC activity in computational model Cumulative distributions for the number of GCs with the indicated percentage of new, lost, or stable spines, respectively. a. For strong dependence of the spine formation on MC activity (*α*0 = 0.1, *α*1 = 1.8) the fraction of stable spines decreased with odor exposure (right panel), whereas the fraction of new and lost spines increased (left and middle panels), contrary to the experimental findings (cf. Fig.1i). b. Without dependence of the formation rate on MC activity (*α*0 = 1.0, *α*1 = 0) the shifts in the distribution functions are still consistent with the experiments. Despite the small shifts, they are highly significant due to the large number of spines available in the simulations.

**Extended Data Figure 11:**
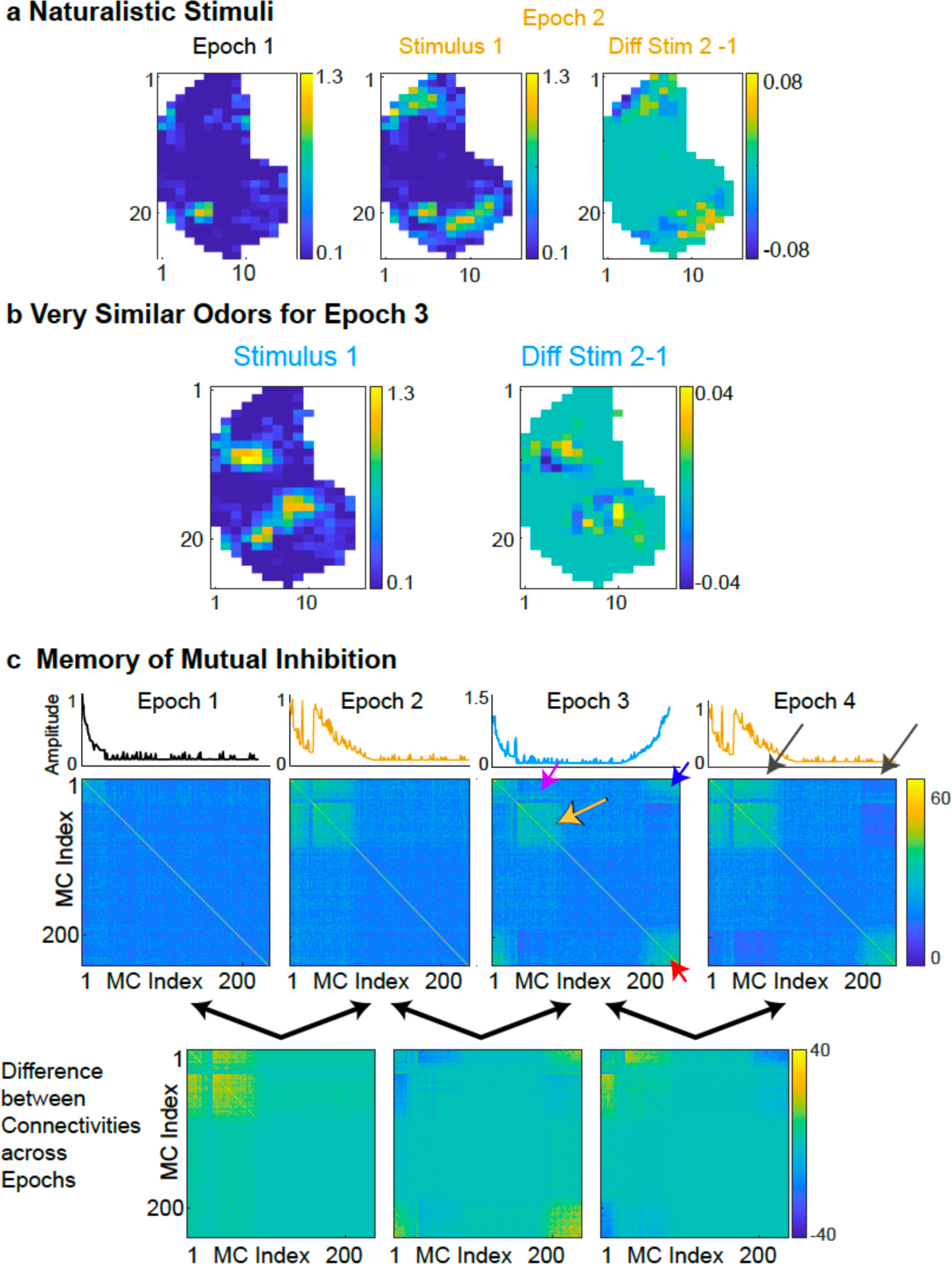
Memory persists in computational model through the presentation of novel odors. a. Naturalistic stimuli for cage odor (butyric acid) and odor mixtures in epoch 2 (isopropylbenzene and cyclo-hexanone). The right panel shows the small difference between the activation patterns of the two mixtures. b. Glomerular activation patterns for 2 novel odors comprised of very similar mixtures of octanal and nonanal. Stimulus 1 is shown in the left panel. The small difference between the two stimuli is shown in the right panel. c. Strength of the disynaptic mutual inhibitory connections between MCs at the end of the four epochs. The MCs are ordered such that the cage odor activated predominantly MCs 1 to 30, the stimuli of epoch 2 drove mostly MCs 30 to 100 and those of epoch 3 MCs 180 to 236, as indicated above the connectivity matrices. The differences in the connectivities are shown in the bottom panels. In epoch 3 new connections reflecting the novel stimuli are added (red arrow) without a substantial reduction in the connections among the MCs activated by the stimuli of epoch 2 (yellow arrow). Connections between MCs activated by the cage odor and the stimuli of epoch 2 were removed (purple arrow), since the associated GCs were activated in the intermediate range in which spines are deconsolidated (yellow arrow in Fig.7b bottom right panel). Re-exposing the network to the stimuli of epoch 2 affected mostly the MCs that were activated by those stimuli as well as the cage odor (gray arrows).

**Extended Data Figure 12:**
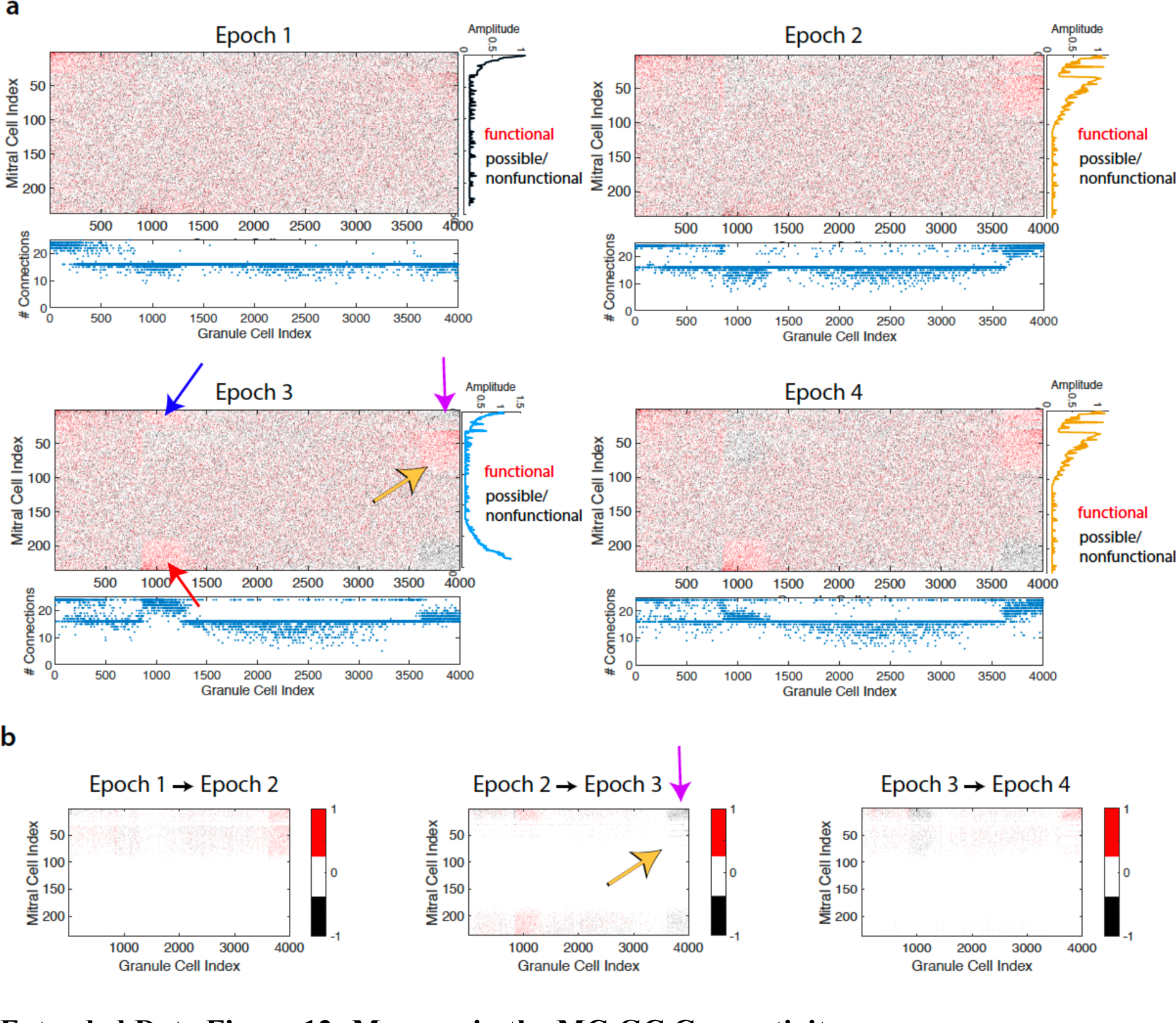
Memory in the MC-GC Connectivity. a. MC-GC connectivities (red dots: functional connections, black dots: nonfunctional spines or possible spine locations) at the end of the four epochs (cf. Extended Data Figure 11). The GC are ordered such that the cage odor activates predominantly GCs 1 to 400, the odors of epoch 2 drives GCs 3500 to 4000, and the odors of epoch 4 activates GCs 800 to 1300. The corresponding MC activity patterns are indicated along the vertical axes. The bottom panels indicate the total number of connections of each GC. Odor stimulation established connections to the activated MCs and increased the total number of connections of the GCs involved. During epoch 3 the GCs that were predominantly activated by the stimuli of epoch 2 preserved the connections to the MCs activated by those stimuli (yellow arrow), but lost connections to MCs predominantly activated by the cage odor (purple arrow). b. Changes in the connectivities between epochs. Red dots indicate new connections, black dots mark lost connections.

**Extended Data Figure 13:**
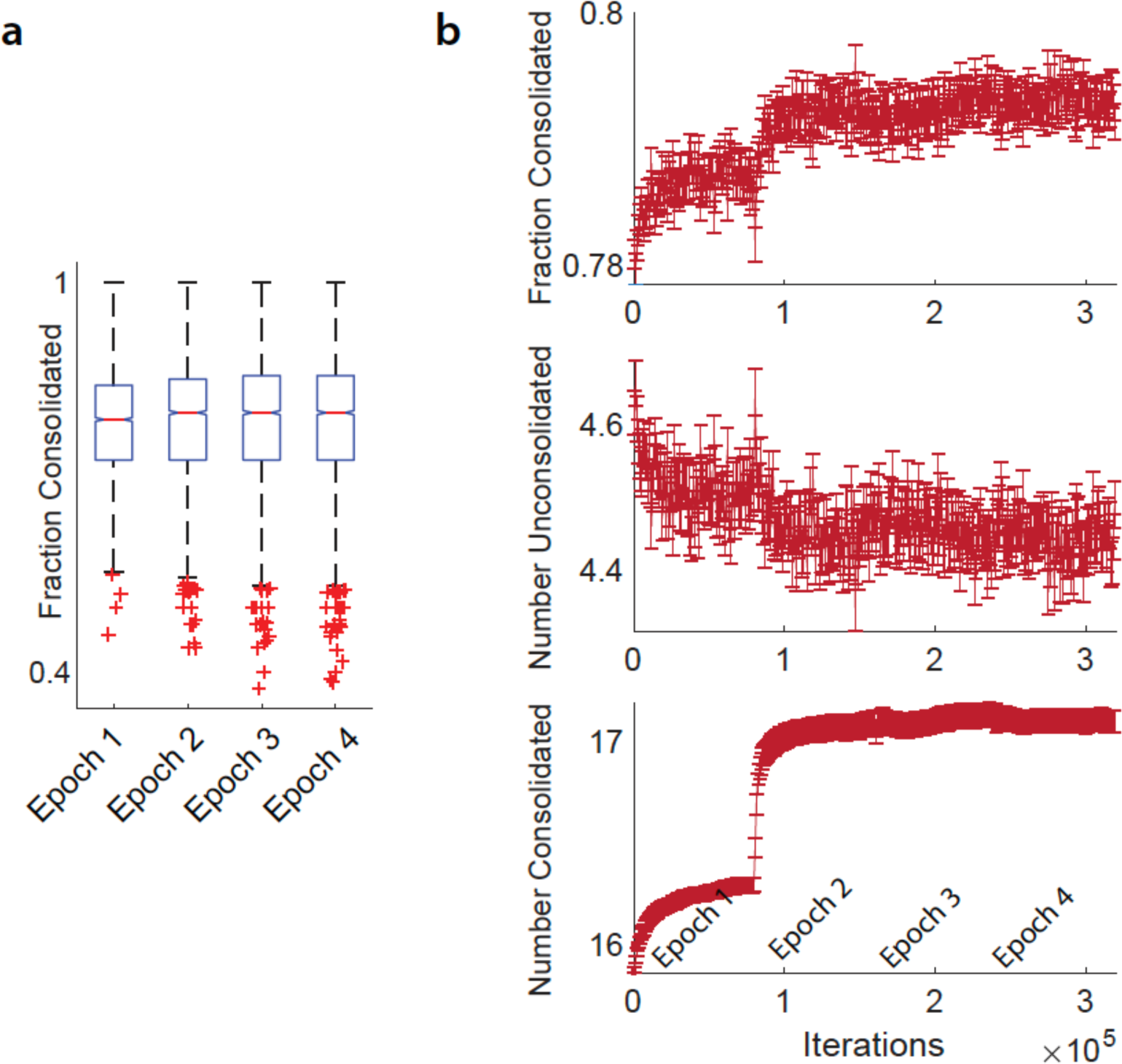
Spine Consolidation during Training The changes in the number of consolidated spines and in the fraction of consolidated spines during learning protocol of Fig.7h are small on average (cf. Extended Data Figure 11). a. Statistics of the fraction of consolidated spines among all spines. The change in the mean from epoch 1 to epoch 2 was significant (p<0.05, the notches do not overlap). Across the subsequent epochs the change in the mean was not significant, even though the connectivities changed significantly (cf. Extended Data Figures 11,12). b. The evolution of the fraction of consolidated spines (top panel) as well as the total number of consolidated and unconsolidated spines (lower 2 panels). The change in the number of consolidated spines is small, but highly significant.

**Extended Data Figure 14:**
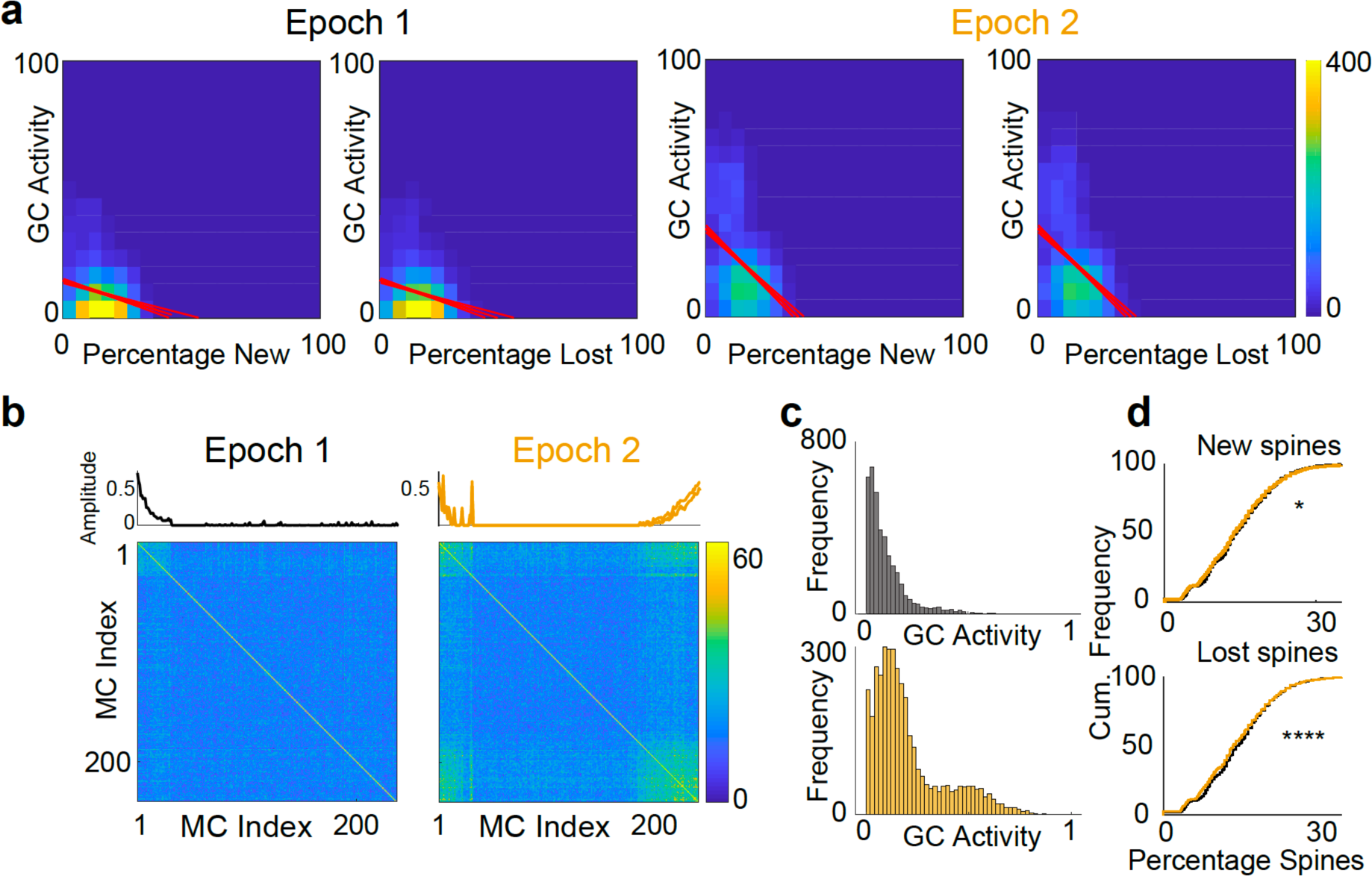
Computational model of spine dynamics, additional data. a. New and lost spine correlation with GC activity, substantially stronger correlation after odor exposure in epoch 2 (cf. Fig.1i). b. Disynaptic MC-MC inhibition. MCs are ordered according to their activation by the stimuli as indicated by the panels above the connectivity matrices. c. GC activity distributions after epoch 1 and 2. d. Stimulus exposure increased the fraction of stable spines and reduced that of new and lost spines. The shifts in the cumulative distributions were small, but highly significant.

